# Pre-Cambrian origin of *envelope*-carrying retrotransposons in metazoans

**DOI:** 10.1101/2023.10.26.564294

**Authors:** Shashank Chary, Rippei Hayashi

## Abstract

Retrotransposons or endogenous retroviruses (ERVs) essentially carry open reading frames of *gag* and *pol*, which are utilized to selfishly replicate themselves in the host germline genome. One rare example of ERVs that additionally carry *envelope* genes is *Ty3/gypsy* errantiviruses in Drosophila. Though they are structurally analogous to retroviruses, it remained unclear whether *envelope*-containing *Ty3/gypsy* elements represent recent, lineage-specific acquisitions of viral fusogens or an ancient association between retrotransposons and *envelope*-like genes. We systematically searched for intact *envelope*-containing ERVs that are homologous to *Ty3/gypsy* in invertebrate metazoan genomes and found that they are widespread across taxa including ancient animals such as cnidarians, ctenophores and tunicates. Many elements occur as multiple highly similar copies in their respective genomes, consistent with recent genomic expansion in some host lineages. *Envelope* genes are classified into those that resemble glycoprotein F from paramyxoviruses and glycoprotein B from herpesviruses, and both types are equally abundant and widespread. Phylogenetic and structural analyses revealed that *envelope* genes have largely diverged with *pol* genes as well as with the host organisms throughout their evolutionary history and recombined infrequently, suggesting that the *envelope* acquisition to ERVs is ancient and likely dates to before the split of bilaterian and non-bilaterian animals in Pre-Cambrian era.

## Introduction

Long terminal repeat (LTR) retrotransposons are one of the most abundant classes of transposons that are ubiquitously present in eukaryotic genomes (Havecker et al. 2004; Wicker et al. 2007). They replicate by reverse-transcribing the messenger RNA and inserting the DNA copy into the host genome (Wilhelm and Wilhelm 2001). They essentially carry two open reading frames (ORFs), namely, *gag* and *pol*. *gag* encodes for a structural protein that packages RNA into capsids while pol encodes a multi-functional protein that contains the reverse transcriptase, RNase H and integrase domains. *Gag* and *pol* ORFs are flanked by 5’ and 3’ LTRs that contain regulatory sequences required for transcription and replication. Reverse transcription is primed from a tRNA primer-binding site located near the 5’ LTR. Although intact copies can continue to replicate, individual insertions acquire mutations and degrade over time once they become inactive. Some retrotransposon remnants remain in the genome to provide functional ORFs and regulatory sequences of host gene expression (Chuong et al. 2017).

LTR retrotransposons are divided into several major lineages, including *Ty1/copia*, *BEL/pao* and *Ty3/gypsy* elements, which are distinguished primarily by POL sequence similarity and domain organisation (Eickbush and Jamburuthugoda 2008). *Ty3/gypsy* retrotransposons are especially closely related to retroviruses among LTR retroelements and typically encode GAG and POL proteins, with POL providing the enzymatic activities required for reverse transcription and integration. They are broadly distributed across eukaryotes, including animals, fungi and plants, although their abundance, domain architecture and ORF arrangement vary substantially between lineages. In this study, we use “*Ty3/gypsy* retrotransposon” for elements whose POL proteins classify with the *Ty3/gypsy* group, as described in the Methods.

Retroviruses are closely related to LTR retrotransposons, but differ from most retrotransposons by carrying an envelope gene, *env*, which encodes a membrane fusion protein essential for infectivity (Xiong and Eickbush 1988; Xiong and Eickbush 1990). Although they typically horizontally transmit between individual animals, they can occasionally infect the germline cells of the host, vertically transmit thereafter, and become known as endogenous retroviruses (ERVs). A well-studied example of such endogenization events is found in Koala, in which nearly identical copies of an exogenous lymphoma-causing retrovirus are found in the genomes of geographically distinct Koala populations (Tarlinton et al. 2006; Zheng et al. 2020). Although individual ERV insertions are subject to genetic drift and degrade over time after they acquire mutations, some insertions remain infectious and continue to make new insertions in the germline genome (Ribet et al. 2008; Miyazawa et al. 2010; Anai et al. 2012; Verneret et al. 2025). However, it is unclear how long such infectious ERVs vertically transmit in a long evolutionary time. When homologous ERVs are found in distantly related species, it can be either seen that they have been inherited from their common ancestor or recently transmitted from one species to another through horizontal transfer. The mechanism of such horizontal transfer events is even less clear although it is often hypothesised that exogenous retroviruses that are borne by one species infect the other species that shares the habitat (Hayward et al. 2020; Simpson et al. 2022; Mottaghinia et al. 2024).

Envelope-like genes are not restricted to vertebrate retroviruses. Canonical vertebrate retroviruses generally encode retroviral class I Env proteins, whereas some ancient retrovirus-like lineages, such as lokiretroviruses, encode Env proteins related to Mononegavirales-like fusion glycoproteins, including paramyxovirus/pneumovirus-like F proteins (Wang and Han 2022). *env*-like ORFs have been reported in several retroelement contexts, including *Ty3/gypsy*, *Ty1/copia*-like elements, and *BEL/Pao* elements such as the Drosophila element *Roo*, and mosquito *BEL/Pao* elements associated with Chuviridae-like glycoproteins (Malik et al. 2000; de la Chaux and Wagner 2009; Dezordi et al. 2020). Their encoded proteins can resemble unrelated viral fusogens. Known *env*-like proteins associated with invertebrate retroelements include glycoprotein F-like proteins, here referred to as F-type ENV, and glycoprotein B-like class III fusogens, here referred to as HSV/gB-type ENV. These proteins are structurally analogous to viral entry proteins, but their evolutionary origins and the extent to which they mediate infectivity in retroelements remain unclear (Hayward 2017).

The best-characterised *env*-containing retroelements in invertebrate genomes are Drosophila *gypsy* elements (Pélisson et al. 1994; Song et al. 1994; Terzian et al. 2001; Pelisson et al. 2002). These so-called errantiviruses behave like bona fide retroviruses, making virus-like particles in the ovarian somatic cells and infecting the germline cells (Song et al. 1997; Leblanc et al. 2000). Therefore, errantiviruses replicate within the genome of the same animal as other retrotransposons do but through an intra-gonadal infection mechanism. Errantiviruses are widespread in Drosophila genomes and form a monophyletic group among other Drosophila *gypsy* elements based on the phylogeny of the *pol* ORF, suggesting that a subtype of *gypsy* acquired the *envelope* gene in the common ancestor of Drosophila and has been vertically transmitting thereafter (Bargues and Lerat 2017). It is postulated that the ancient *gypsy* retrotransposon in Drosophila acquired the *envelope* gene from an insect-specific DNA virus, baculovirus, through recombination (Malik et al. 2000; Rohrmann and Karplus 2001). However, *Ty3/gypsy* LTR retrotransposons with *envelope* genes have also been sporadically documented in non-insect animals, such as in nematodes, rotifers, and tunicates, obscuring their true origin (Volff et al. 2004; Rodriguez et al. 2017; Sacco et al. 2022).

Here we comprehensively searched for *envelope*-carrying *Ty3/gypsy* retrotransposons in invertebrate metazoan genomes. We found elements with uninterrupted ORFs flanked by putative LTR sequences in nearly every major phylum, including ancient animals such as cnidarians, ctenophores, and tunicates. Furthermore, many of these elements occur as multiple highly similar copies in their respective genomes, consistent with recent or ongoing genomic expansion in some host lineages. Phylogenetic and structural analyses of the *pol* and *envelope* genes show that they have co-evolved with their host species for a long evolutionary time. Therefore, we propose that the acquisition of the *envelope* gene to retrotransposons occurred anciently in metazoans, and they have been vertically transmitting since pre-Cambrian era.

## Results

The approach to study the phylogeny and evolution of LTR retrotransposons, is conventionally to mine the genome for sequences which share homology to retroviral *gag*, *pol*, or *envelope (env)* genes. Prior studies on ERVs and retrotransposons have assembled trees based on reverse transcriptase (RT) or Integrase (IN) from *pol* to obtain insights (Xiong and Eickbush 1990; Wang and Han 2021). One can also investigate synteny of loci between different hosts, estimating an age of origin for those elements (Bonnaud et al. 2005; Ueda et al. 2020).

This study aims to investigate the evolution of *env* genes acquired by intact *Ty3/gypsy* retrotransposons, and to explore structural variations within those elements. The conventional approach, therefore, needs to be altered to approach this. We only consider intact copies that retain uninterrupted open reading frames (ORFs) for GAG, POL and ENV, because non-degraded ORFs provide more reliable material for domain annotation, structural prediction and phylogenetic comparison (Fig 1A). Because intact ORF structure was part of our search strategy, the recovery of intact ORFs should not by itself be interpreted as evidence of recent mobilisation. Where possible, we also recorded whether closely related copies were present in the same genome, because multicopy structure provides additional evidence consistent with recent or ongoing genomic expansion. The study of syntenic loci here is inappropriate, as the genomic location of recently mobilised transposons will likely never be the same in evolutionarily distant hosts. Like most studies of repeat elements, the sequences are highly diverse, and we therefore supplement phylogenetic evidence with structural predictions.

**Figure 1.**
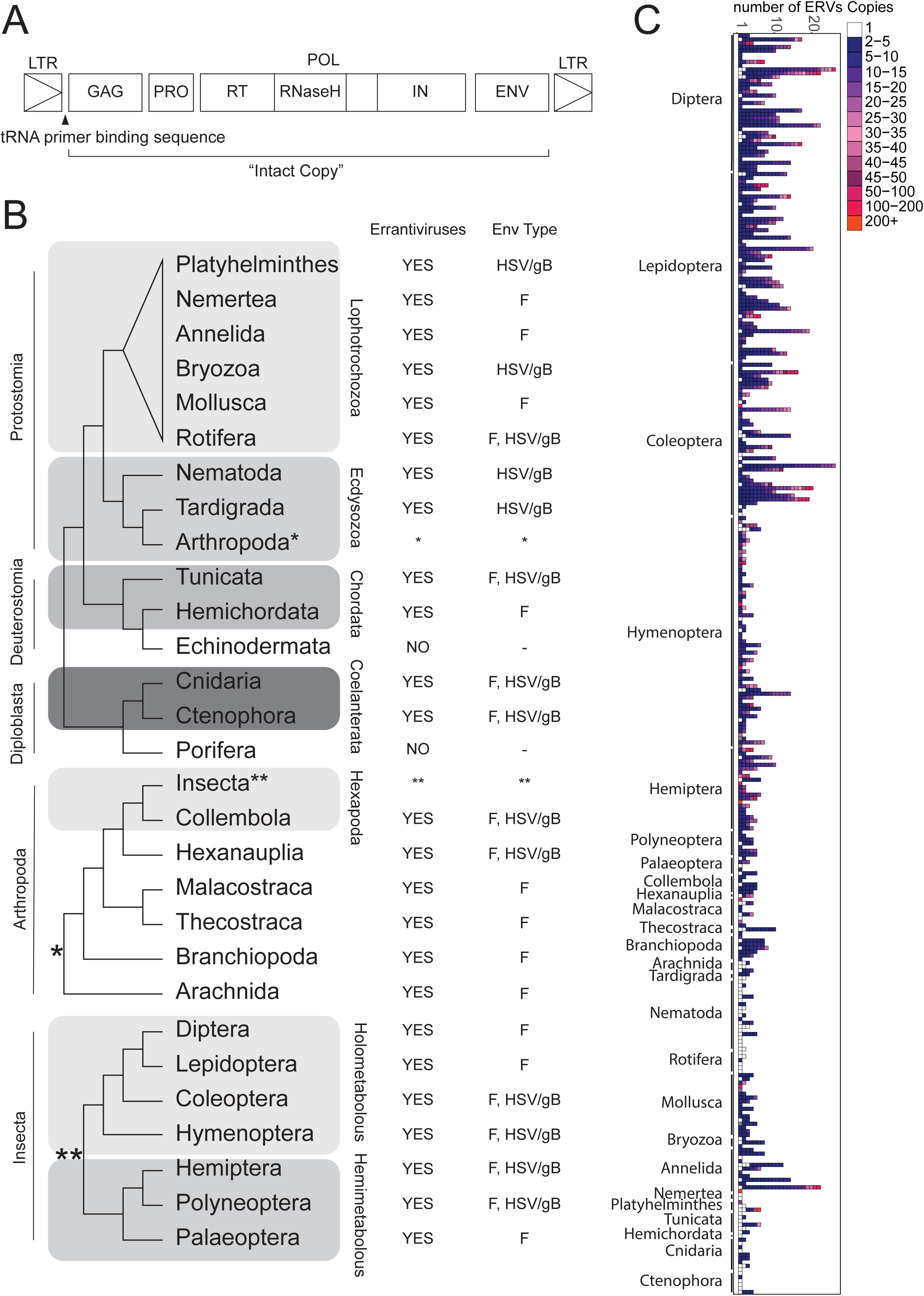
– The widespread nature of intact GAG-POL-ENV elements across metazoan genomes. **(A)** Open reading frame arrangement of intact errantivirus copies. GAG encodes the structural capsid-like protein, POL encodes the enzymatic protein containing protease (PRO), reverse transcriptase (RT), RNase H and Integrase (IN) domains, and ENV encodes the envelope-like fusogen. tRNA primer binding sequence is used for reverse transcription. **(B)** Summary of Errantiviruses found in Metazoan phyla with type of ENV protein indicated. **(C)** Copy number of intact errantivirus copies in metazoans. Column represents a genome assembly, each tile representative of annotated errantivirus, colour of tile represents copy number.

We systematically searched for homologous elements of Dipteran *Ty3/gypsy* errantiviruses in the referenced genomes across metazoan phyla using the annotated protein coding sequences of *gag*, *pol* and *env* for tBLASTn searches (see methods and Fig. S1). The first tBLASTn search identified elements outside Diptera, therefore, we extended the search using the representative of newly identified elements as baits. We repeated this process and examined whether the additional baits uncovered elements that were not represented in the previous tBLASTn searches by identifying new clades in the phylogenetic tree of POL (see methods and Fig. S1). After five iteration of tBLASTn searches, we no longer discovered elements that formed a new POL phylogenetic clade (Fig. S1B). These steps allowed us to obtain a comprehensive set of elements in metazoan genomes that are homologous to Dipteran *Ty3/gypsy* errantiviruses (Table S1, summarised in Fig 1). The majority of the newly identified genomic copies carry putative LTRs and tRNA primer binding sequences (1151 out of 1512 elements, Table S1), suggesting their recent integrations. Additionally, most of the annotated elements sit within contigs larger than 0.1 mega bases (1434 elements, Table S1), indicating their true origins in their respective genomes. We generated maximum-likelihood phylogenetic trees from multiple sequence alignments of two conserved Pol regions: the Integrase (IN) domain and an extended RT/connection region. The extended RT/connection region includes the RT polymerase core and the downstream connection subdomain, and is therefore broader than the RT polymerase core alone but does not include the complete canonical RNase H domain analysed structurally in Figure 6. ORFs of annotated Drosophila *gypsy* errantiviruses were also included in the trees to aid classifications.

We performed a similar analysis on vertebrate genomes using known vertebrate retroviral *gag*, *pol*, and *env* sequences as bait in addition to Drosophila errantiviruses. We did not identify any intact ERVs in vertebrate genomes that were homologous to invertebrate elements, nor did we find elements in invertebrate genomes that predicted homology to vertebrate elements. There is therefore a clear phylogenetic separation between invertebrate elements and vertebrate ERVs, despite structural similarities. The investigation on the vertebrate ERVs will be reported elsewhere, as it is outside the scope of the present study.

For clarity, we use the following terminology in this study. ERV refers to an endogenous retrovirus or retrovirus-like LTR retrotransposon present in the host genome. *Ty3/gypsy* retrotransposon refers to an LTR retrotransposon classified with the *Ty3/gypsy* group based on POL similarity (see methods). Errantivirus is used provisionally and operationally to refer to *env*-containing *Ty3/gypsy* retrotransposons identified in this study, following the historical use of this term for *env*-containing *Metaviridae/Ty3/gypsy* elements (Boeke et al. 2000). This usage is not intended as a formal taxonomic proposal and does not imply that all such elements share the experimentally characterised Drosophila *gypsy* lifecycle. We use lower-case *env* for the gene and upper-case ENV for the encoded protein. F-type *env*/ENV refers to glycoprotein F-like *env* genes or proteins, whereas HSV/gB-type *env*/ENV refers to herpesvirus glycoprotein B-like *env* genes or proteins.

### Multiple intact elements of *env*-carrying *Ty3/gypsy* retrotransposons are found widespread across metazoan species

We found that intact copies of errantiviruses carrying all three retroviral ORFs are found not only outside of insect orders but are ubiquitously present across most major metazoan phyla. This includes species from ancient phyla such as Cnidaria, Tunicates and Ctenophora, but excludes Echinodermata and Porifera (Fig 1B and Table S1). To test whether the apparent absence of env-carrying Ty3/gypsy elements in Porifera and Echinodermata could be explained by poor assembly quality or a general failure to recover full-length retrotransposons, we searched two representative genomes from each phylum for full-length GAG-POL *Ty3/gypsy* elements using the same general annotation strategy. We identified multiple full-length, multicopy GAG-POL elements in all four genomes examined, many of which were flanked by predicted LTR sequences and associated with putative tRNA primer-binding sites (Table S3). Thus, the lack of detectable GAG-POL-ENV elements in these representative Porifera and Echinodermata genomes is unlikely to be explained simply by an inability to recover intact *Ty3/gypsy*-like retrotransposons from these assemblies. Many carry ENV proteins that are homologous to glycoprotein F, commonly found in paramyxoviruses, such as Nipah, Hendra and Respiratory Syncytial Viruses (McLellan et al. 2013; Dang et al. 2019), but is also carried in insect baculoviruses (and Drosophila errantiviruses) (Malik et al. 2000; Rohrmann and Karplus 2001). Other elements we identified instead carry ENV that resembles glycoprotein B (gB), famously found as part of the glycoprotein complex in Herpes Simplex Viruses (HSV). Both types of ENV glycoproteins are carried by errantiviruses found in ancient hosts such as rotifers, tunicates, cnidarians and ctenophores, as well as in hosts from insect and arthropod orders (Fig 1B).

Though, errantiviruses found in Nematoda, Bryozoa and Platyhelminthes only carry HSV/gB-type ENV, and the ORFs are arranged as GAG-ENV-POL rather than GAG-POL-ENV. These GAG-ENV-POL elements do not form a single monophyletic group in the POL phylogenetic tree, but instead occur in distinct host-associated clades, as shown later in Supplementary Figure S4B and Supplementary Figure S8. Most elements that were identified have multiple intact copies in their respective genomes (as defined as copies with >98% nucleotide identity across >98% of the three ORFs, see methods, Table S1), a hallmark feature of recent transposition (Cordaux and Batzer 2009; Gagnier et al. 2019), indicating that most elements that we have annotated likely behave as bona-fide retrotransposons (Fig 1C). Activity of these elements are highly varied, as the copy number can be as little as one intact copy, to hundreds of intact copies in the genome (Fig 1C, Table S1). Importantly, multi-copy ERVs are present in nearly all animal phyla where we see intact elements, suggesting that these elements have been recently transposing in a wide range of metazoan species.

The phylogenetic tree assembled on the intact errantivirus *pol* genes shows congruence, meaning that the tree assembled on extended RT/connection region closely reflects that of IN. The simplified tree in Figure 2 is used to summarise the major phylogenetic groups; the complete POL extended RT/connection tree, the corresponding IN tree and their congruence are shown in Supplementary Figure S2. Errantiviruses within most clades in one tree are represented in the same or closely related clades in the other (Fig. S2B). Importantly, the errantiviruses found in phyla outside of insects, are in a different clade to those found within, which we call “ancient errantiviruses” and “insect errantiviruses”, respectively (Fig 2 and S2A). Notably however, few elements found in insects sit within clades of ancient errantiviruses, rather than other insect ones (Fig 2). Annotated Drosophila *gypsy* elements included in the tree, namely, Gypsy-ZAM, Gypsy-Gypsy/Gypsy-5_DGri and Gypsy-DM176/Idefix present themselves in different clades while they cluster together with other insect errantiviruses, in an arrangement with previous studies that have classified Drosophila *gypsy* elements (Bargues and Lerat 2017) (Fig 2). Though, one element from a cnidarian species *Paleozoanthus reticulatus* clusters together with insect elements, in a clade adjacent to DM176 and Idefix (Fig 2), its origin is unclear as the element is found on a short genomic contig (Table S1).

**Figure 2.**
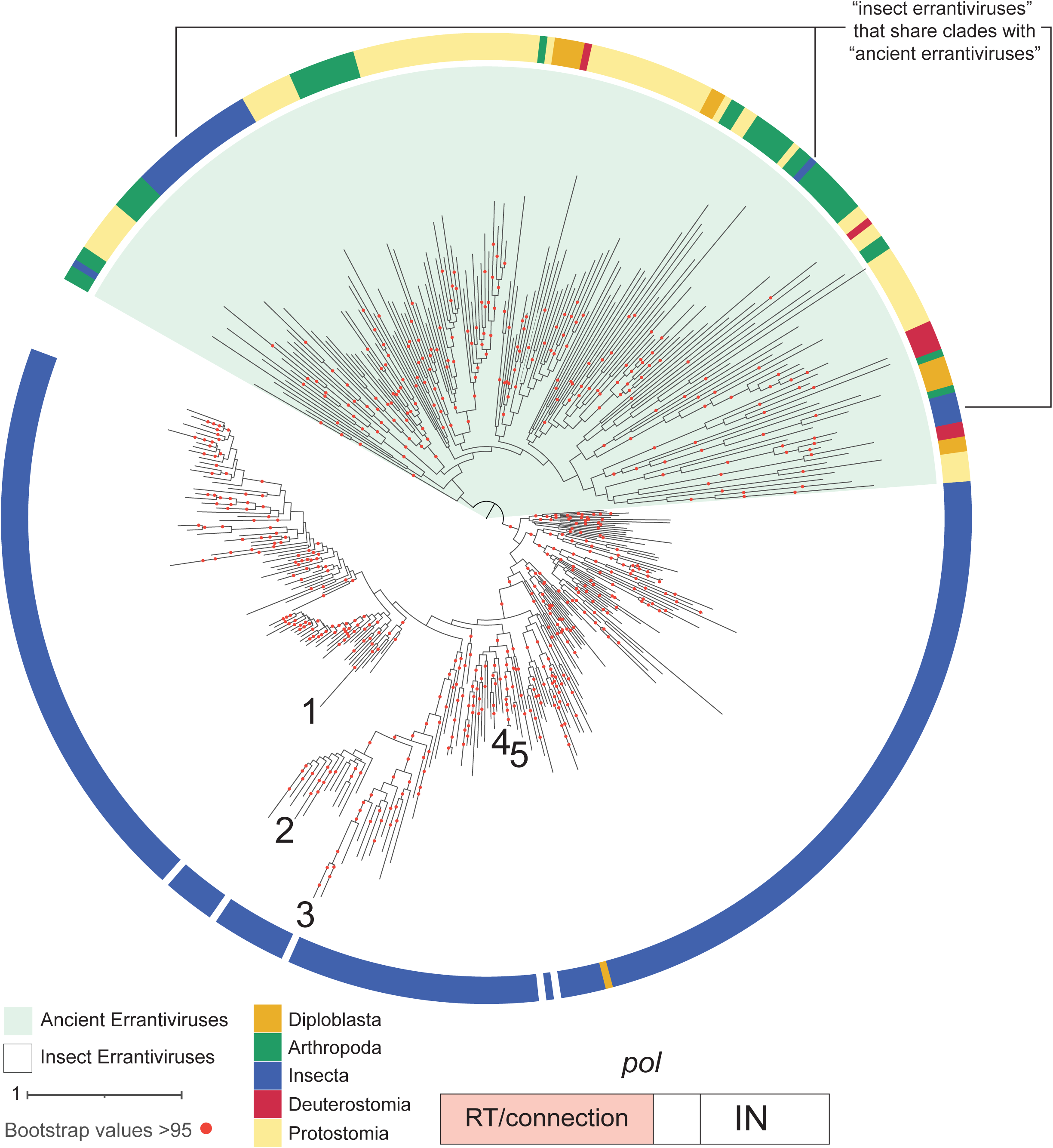
– Phylogeny of POL extended RT/connection region from intact metazoan errantiviruses. Maximum likelihood tree constructed on representative POL extended RT/connection regions of intact errantiviruses (see methods) and rooted to the midpoint. The “ancient errantiviruses” and “insect errantiviruses” phylogenetic groups are highlighted in green and white respectively. Positions of known dipteran errantiviruses, Gypsy_ZAM (1), Gypsy_Gypsy5_Dgri (2), Gypsy_Gypsy (3), Gypsy_Idefix (4), and Gypsy_DM176 (5) are indicated. Host classifications of errantiviruses are annotated in colour, non-arthropod Protostomia (yellow), Deuterostomia (red), Diploblasta (orange), non-insect Arthropoda (green), Insecta (blue). One cnidarian element within the group of insect errantiviruses is marked by an asterisk.

Many of the newly identified “insect errantiviruses” sit in separate clades to the Annotated Drosophila *gypsy* elements (Fig 2, tree positions I3 – I6 in Fig. S2A). These “insect errantivirus” clades are strongly supported by bootstrap values greater than 95 and are additionally supported by the annotations of the tRNA primer binding site (PBS) found in each element. For example, Tree position I1 within “insect errantiviruses” mainly carry tRNA^Ser^, whereas Tree position I2, consisting of only dipteran elements, predominantly use tRNA^Lys^, though few use tRNA^Ser^ or tRNA^Pro^ (Fig. S2A and S2C). Elements in insects within positions I4, I5 and I6 predominantly carry tRNA^Leu^ primer binding sites, in addition to Serine and other uncommon tRNAs (Fig. S2C). This is mostly consistent with past works, that have identified Drosophila *gypsy* elements to carry either Serine or Lysine tRNA PBS (Terzian et al. 2001; Stefanov et al. 2012). Notably, most discrete clades comprised elements from the same insect orders, suggesting long term vertical transmission and divergence within related species (Fig. S3). A detailed view of the insect errantivirus POL extended RT/connection clades, including host order, ENV type and the positions of phylogenetic groups I1–I6, is provided in Supplementary Figure S3.

The “ancient errantivirus” group also consists of clades strongly supported by bootstrap values greater than 95, though the tRNA PBS content within the clades is much more divergent (Fig. S4C). Notably however, errantiviruses found in insects that sit within this group, in tree positions A1, A2 and A7 continue to carry predominantly tRNA^Ser^, tRNA^Lys^ and tRNA^Leu^ (Fig. S4B). Discrete clades are consistently made up of elements in hosts from the same, or closely related phyla (Fig. S4B), for example, tree position A5 consists of elements found in Thecostraca, Annelida and Mollusca, each of which form their own discrete clade (Fig. S4B). This indicates that they were either present in their common ancestor or inherited more recently within the same animal group. Given that neighbouring clades consist of elements found in related animal phyla, it is more likely that they are ancient in nature. For example, clades that consist of insect elements are flanked by clades of other arthropod elements (Fig. S4B, tree position A2). Clades containing Annelida elements often cluster with or next to mollusc elements (Fig. S4B, tree positions A5-A7). This congruence between clades and species is highly indicative of long-term vertical transfer, as known in non *env-*containing retrotransposons (Sormacheva et al. 2012; Wanner and Faulk 2021). Within larger clades containing a wider variety of host orders such as tree position A9, the separation is still evident and strongly supported clades only consist of elements from the same order of host (Fig. S4B). Supplementary Figure S4 provides the full phylogenetic context for the “ancient errantivirus” group, including host taxonomy, ENV type, tree positions A1–A11 and tRNA PBS annotations.

To test the host-taxonomic structure more directly, we performed targeted permutation analyses in two well-sampled POL extended RT/connection subclades in tree positions I1 (Fig. S3) and A6 (Fig. S4B), consisting of elements from Lepidoptera and Annelida, respectively (Fig. S5). We scored internal nodes with bootstrap support of ≥95 and 3–10 descendant tips, then randomly permuted host-taxonomic labels across the fixed tree topology while preserving the observed label counts. In the Annelida clade, supported small subclades were far more host-concordant than expected by chance: all six scored subclades consisted of elements from the same host family, compared with a null expectation of 0.27 subclades, and four of six consisted of elements from the same host species, compared with a null expectation of 0.13 subclades (10,000 permutations; empirical P = 1.0 × 10^-4^ for both metrics). The Lepidoptera clade showed a more mixed pattern, but several highly supported subclades were enriched for related host groups at the superfamily or broader taxonomic level. These targeted analyses do not exclude rare horizontal transfer, particularly between closely related hosts, but indicate that host labels are not randomly distributed across local POL phylogenetic topologies; the analysed Annelida A6 and Lepidoptera I1 subclades are shown in Supplementary Figure S5.

Furthermore, clades formed on the trees using *pol* proteins tend to carry the same type of *env* (Fig 3A and S4A). For example, within A6 in the “ancient errantivirus” group, elements that carry HSV/gB-type *env* are in a separate clade to those carrying F-type *env* (Fig. S4B). Though some clades appear to be a mix of both, elements with HSV/gB-type *env* still tend to cluster together, indicating that they have stayed with *pol* for long evolutionary periods of time.

**Figure 3.**
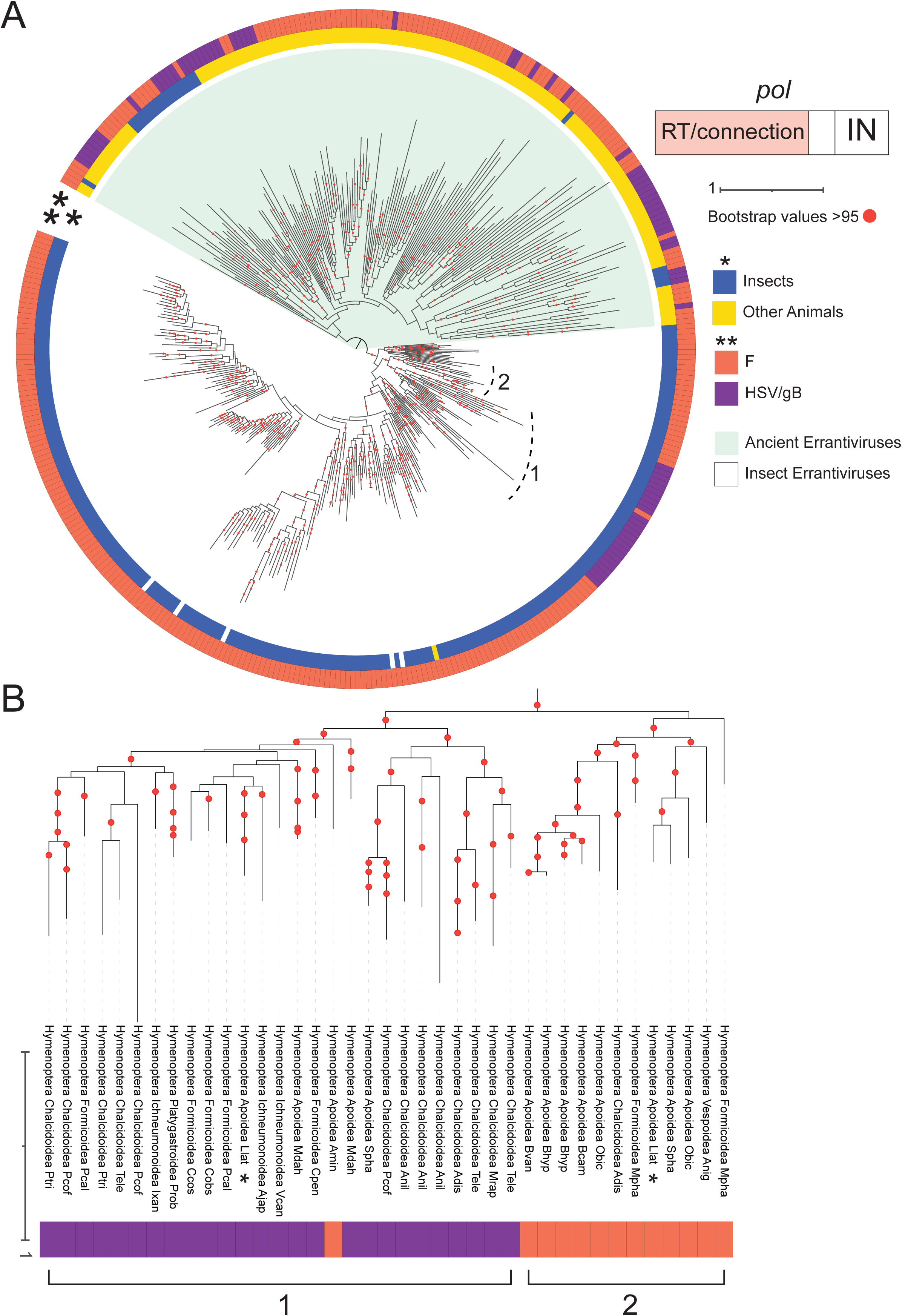
– *env* classifications of intact metazoan errantiviruses. **(A)** Shown is a POL extended RT/connection maximum likelihood tree of the representative errantiviruses (see methods) with host (*) and type of ENV protein (**) indicated. Colours represent non-insect animals (*) (yellow), insects (*) (blue), glycoprotein F (F) (**) (orange), and glycoprotein B (HSV/gB) (**) (purple). Clades containing hymenopteran elements in “Insect errantiviruses” that carry either F or HSV/gB *env* are annotated as 1 and 2. **(B)** Close up of clades 1 and 2 in “Insect errantiviruses”. F-type and HSV/gB-type errantiviruses that are both found in *Lasioglossum lativentre* are marked by asterisks.

### ENV glycoproteins carried by errantiviruses are highly structured, and their divergence follows that of *pol*

As previously mentioned, we found that errantiviruses can carry HSV/gB-type or glycoprotein F-type *env*. While most HSV/gB-type *env* is found in the “ancient errantivirus” clade, one clade in “insect errantiviruses” exclusively consists of hymenopteran errantiviruses which carry both HSV/gB-type and F-type *env* genes (Fig 3A-B). Interestingly, elements with HSV/gB-type *env* and F-type *env* separate at the base of this clade while they both are found in a wide range of species superfamily within Hymenoptera, suggesting that both HSV/gB-type and F-type *env* have been together with their *pol* genes at least throughout the evolution of hymenopteran species (Fig 3B).

To further characterise the ENV proteins carried by errantiviruses, we identified the ectodomains within the *env* ORF, which is flanked by signal peptides and a transmembrane domain, using DeepTMHMM (https://doi.org/10.1101/2022.04.08.487609). We then analysed trimeric configurations of these ectodomains from representative errantiviruses from the phylogenetic tree using Alphafold 2 (Jumper et al. 2021).

F glycoproteins analysed are highly structured and consistently form four domains that are highly conserved across all elements (Fig 4A). They also predicted a highly conserved Furin cleavage site and hydrophobic fusion peptides in the ectodomain (Fig 4A and S6). The alignment-level evidence supporting conservation of the predicted furin cleavage site and hydrophobic fusion peptide is shown in Supplementary Figure S6. The 3D fold of the ectodomain structures forms a “head” (domains 1-3) and a “stalk” (domain 4), strongly resembling 3D structures of F ectodomains characterised in previous studies, such as those from RSVs (McLellan et al. 2013) (Fig 4C). Most errantiviruses also have an α helix at the C terminus of the ectodomain, referred to as “Domain A” (Fig 4A and 4C). The predicted 3D trimeric structure also reveals two potential configurations of the glycoprotein, based on domain 4: either a long α helix, or two shorter helices. These long and short barrel structures resemble pre– and post-fusion structures of glycoprotein F described in prior studies (McLellan et al. 2013). These highly structured domains in the intact *env* ORFs in these errantiviruses demonstrates that *env* has remained intact and could suggest that their function also remains intact as well.

**Figure 4.**
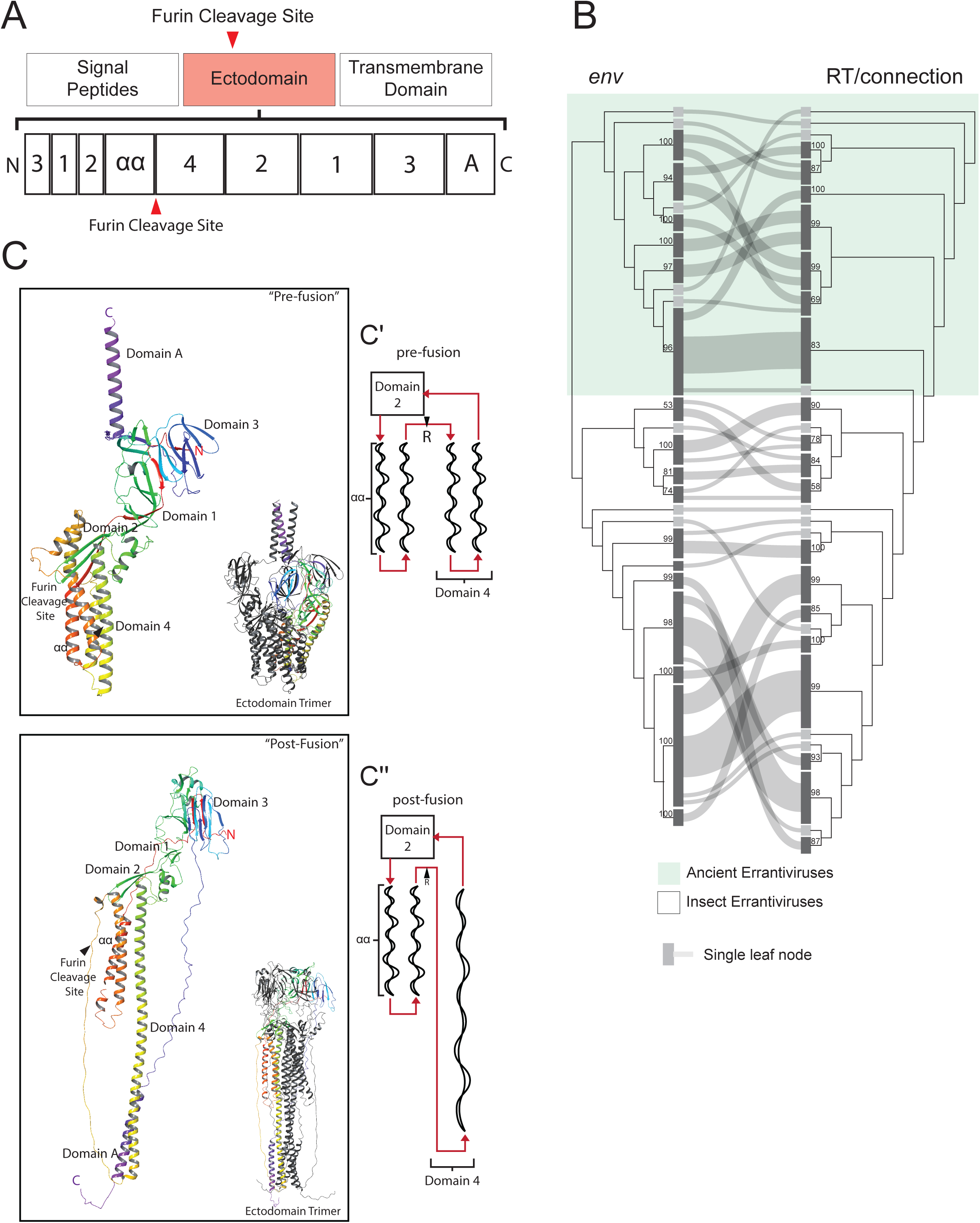
– Structure and phylogeny of glycoprotein F-type ENV found in intact errantiviruses. **(A)** Domain arrangement of glycoprotein F ectodomain in Errantiviruses. **(B)** A Sankey plot of POL extended RT/connection and glycoprotein F-type ENV ectodomain maximum likelihood trees for representative errantiviruses. In the Sankey plot, each ribbon links the clade assignment of the same element in the Pol extended RT/connection tree and the F-type ENV ectodomain tree. The plot is intended as a qualitative summary of global congruence between the two trees. Ribbon crossing can result from the relative ordering or rotation of clades in the two tree drawings and should not by itself be interpreted as recombination. Instead, broad one-to-one mapping between Pol-defined and ENV-defined clades indicates concordance, whereas extensive splitting or many-to-many connections indicates discordance consistent with recombination, env exchange or differential phylogenetic resolution. Bootstrap values for nodes are given. The “ancient errantivirus” group is highlighted in green. **(C)** Monomeric and trimeric structures of the “pre-fusion configuration” and the “post fusion configuration”. Schematic cartoons of F-type ENV ectodomain domain 4 are shown in **(C’)** and **(C’’)** Respectively. Structures are taken from Diptera_Sciomyzidae_Cmar_errantivirus_19 and Branchipoda_Diplostraca_Dcar_errantivirus_3 for “pre fusion” and “post fusion” structures in **(C)**.

To explore the phylogenetic relationships between *pol* and *env*-F, we generated multiple sequence alignments of the ectodomains and POL extended RT/connection regions from representative elements and assembled phylogenetic trees. We found that the ENV ectodomains cluster similarly to the POL extended RT/connection regions, and ENV proteins tend to form clades within the same or closely related species as they do in the POL tree (Fig 4B). Importantly, the ENV ectodomains found in “ancient errantiviruses” cluster separately to those found in “insect errantiviruses”, including those insect elements that present themselves with ancient errantiviruses in the POL tree (Fig 4B). This suggests that *env*-F genes have stayed with *pol*, and the acquisition of *env* likely occurred anciently.

To further compare errantivirus F-type ENV proteins with related fusogens outside *Ty3/gypsy* retrotransposons, we repeated the F-type ENV ectodomain phylogeny after adding representative viral and retroelement F-like proteins, including baculovirus F proteins, paramyxovirus and pneumovirus F proteins, and the *BEL/Pao*-associated Drosophila Roo F-like protein (Fig. S7). The added viral sequences formed family-level clades within the broader diversity of F-type ENV proteins. The inclusion of these external homologues reduced the apparent separation between insect and ancient errantivirus F-type ENV proteins, indicating limited resolution for some deep relationships in the F-type ENV tree. Nevertheless, the errantivirus F-type ENV proteins did not collapse into a shallow *Ty3/gypsy*-specific group, nor did they cluster as a recent derivative of a single sampled viral family. Instead, their diversity was comparable in scale to that separating major viral F-protein groups, consistent with deep diversification of F-type ENV proteins associated with errantiviruses.

The ectodomain of errantiviruses that carry HSV/gB-type *env* are either flanked by signal peptides and a transmembrane domain, or by two transmembrane domains and the Furin cleavage site is not always conserved (Fig 5A). Like F, the trimeric structures of HSV/gB-type ENV in errantiviruses are also highly structured. They carry five highly conserved and distinct domains (designated as domain I – V) arranged in a hairpin configuration, that fold into a trimeric structure with a base, stalk and crown (Fig 5B). Additionally, domain I carries highly conserved “fusion loops” consisting of hydrophobic amino acid residues. These domains have also been characterized in Herpes Simplex Viruses in prior studies (Heldwein et al. 2006; Connolly et al. 2011). Notably, errantiviruses found in insect orders that carry HSV/gB-type *env*, only carry one of these fusion loops (Fig. S8). The domain I–II alignments underlying these cysteine-bridge classifications are shown by host group in Supplementary Figure S8, allowing conservation and variation of cysteine positions to be compared across HSV/gB-type ENV lineages. The structure of the HSV/gB-type ENV carried by these errantiviruses are stabilized by disulphide bonds between cysteine residues, which we refer to as cysteine bridges (Lopper and Compton 2002; Vollmer et al. 2020). These bridges are highly conserved but appear to vary the most within domains I and II of the HSV/gB ectodomain (Fig 5C and S8). Additionally, in some cases, a cysteine bridge within domain I is located near the fusion loops (Fig 5C and S8).

**Figure 5.**
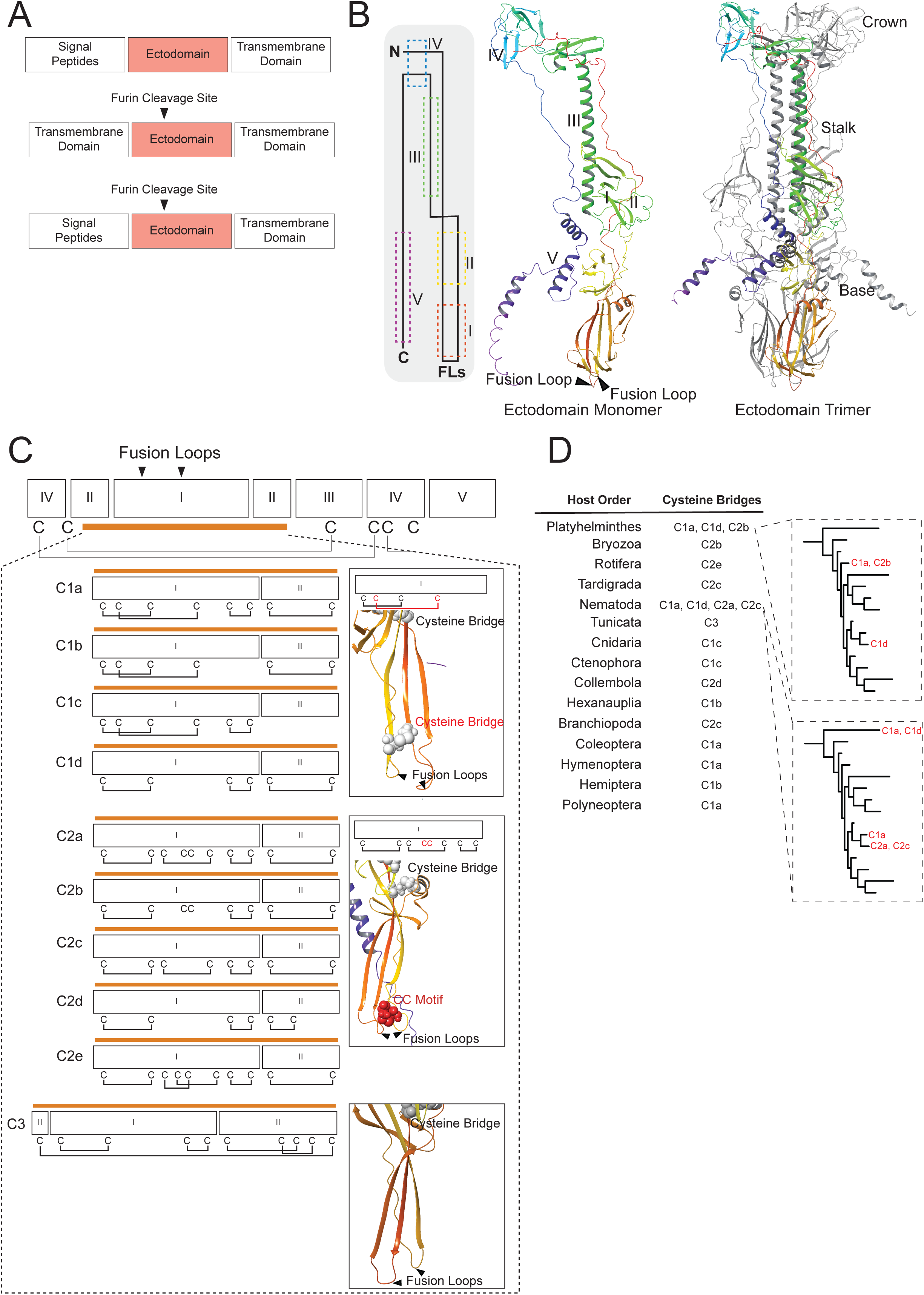
– Structure of glycoprotein HSV/gB-type ENV found in intact errantiviruses. **(A)** Structural arrangements of Ectodomains, Transmembrane domains and signal peptides found in HSV/gB-type errantiviruses. **(B)** Shown are the arrangement of domains of the HSV/gB-type ENV ectodomain, and Alphafold 2-predicted 3D monomer and trimer structures of the ectodomain. The structure is taken from Hemiptera_Pemphigidae_Elan_errantivirus_1. **(C)** Fusion loops and conserved cysteine bridges on structure of the errantivirus HSV/gB-type ENV ectodomain. Shown are variable cysteine bridges in the errantivirus HSV/gB-type ENV ectodomains, in domain I and II (Indicated with orange bar. Proximity of cysteine bridge to fusion loops of domain I are indicated for structures within C1, C2 and C3. Structures taken from Cnidaria_Hydrozoa_Chem_errantivirus_5, Bryozoa_Gymnolaemata_Cpal_errantivirus_2, and Tunicata_Ascidiacea_Clep_errantivirus_1, for C1, C2 and C3 configurations, respectively. **(D)** Shown is the summary of the cysteine bridge configurations found in HSV/gB-type ENV of different metazoan orders. Positions for cysteine bridge configurations within Nematoda and Platyhelminthes shown on Simple schematic. Exact positions of all cysteine bridges can be found in Supplementary Figure S8.

**Figure 6.**
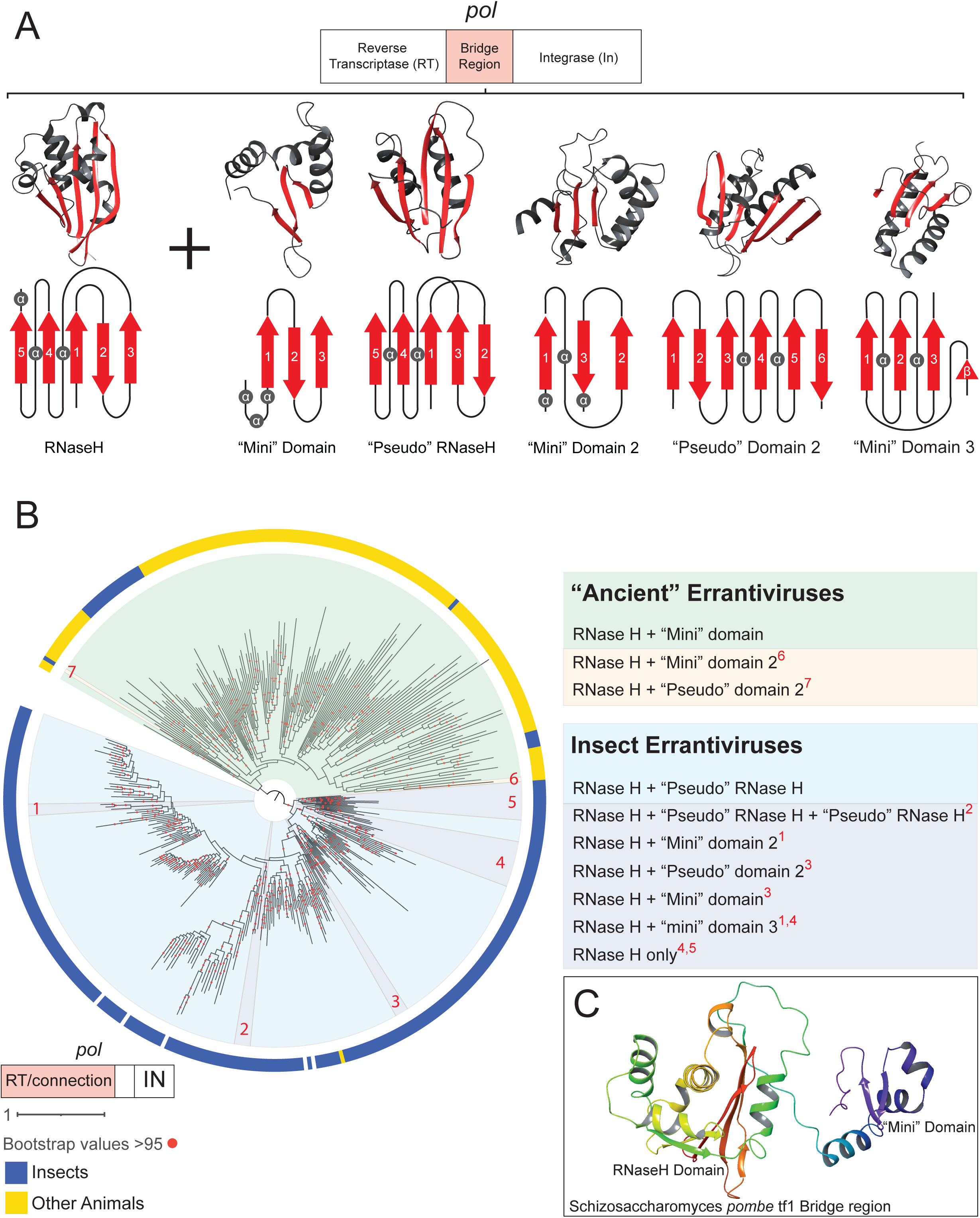
– RNase H domains found in the bridge region of intact errantiviruses. **(A)** Alphafold 2-predicted structures of domains found in the bridge regions of errantiviruses, with topology maps of α helices (grey) and β sheets (red). Structures are taken from Annelida_Polychaeta_Apac_errantivirus_7, Bryozoa_Gymnolaemata_Cpal_errantivirus_3, Coleoptera_Coccinellidae_Npum_errantivirus_1, Annelida_Polychaeta_Slim_errantivirus_9, Diptera_Cecidomyiidae_Smos_errantivirus_1 and Hymenoptera_Apoidea_Obic_errantivirus_2 for RNase H/Tether, ‘mini’ RNase H, ‘pseudo’ RNase H domains, “Mini” domain 2, “Pseudo” domain 2 and “Pseudo” domain 3, respectively. **(B)** Shown is the POL extended RT/connection maximum likelihood tree of representative errantiviruses, with RNase H domain architecture in bridge region of *pol* highlighted. “Insect errantiviruses” with RNase H + pseudo–RNase H (blue), “insect errantiviruses” with bridge region outliers (dark blue), “ancient errantiviruses” with RNase H + mini domain (green), “ancient errantiviruses” with bridge region outliers (yellow). **(C)** The structure of the bridge region of yeast *Schizosaccharomyces pombe Ty3* element *tf1* predicted by Alphafold 2.

To determine whether conservation of cysteine-bridge architecture simply reflected high primary-sequence similarity, we compared errantivirus HSV/gB-type ENV proteins with representative viral and retroelement-associated class III fusogens, including herpesvirus gB, rhabdovirus G, orthomyxovirus GP75/GP64-like proteins, baculovirus GP64 and BEL/Pao-associated gB-like proteins (Backovic and Jardetzky 2009) (Fig. S8). We did not infer a global phylogeny for these proteins because their primary sequences and domain organisations were too divergent for reliable full-ectodomain alignment. Instead, we compared predicted domain organisation, cysteine-bridge architecture and pairwise amino-acid identity within each viral family or errantivirus ENV group.

This comparison showed that conserved cysteine-bridge architecture can persist despite extensive primary-sequence divergence. For example, the sampled herpesvirus gB proteins, representing alpha-, beta– and gammaherpesvirus homologues, shared only 27.8–30.3% pairwise amino-acid identity across aligned residue–residue positions while sharing an identical cysteine-bridge architecture (Burke and Heldwein 2015) (Fig. S8 and Table S4). Rhabdovirus G proteins and orthomyxovirus GP75/GP64-like proteins showed similarly low pairwise identities, 21.8–27.7% and 24.3–30.1%, respectively while also retaining mostly shared cysteine-bridges (Belot et al. 2020). For clarity, we use labels such as C1a and C2b to denote recurrent HSV/gB-type cysteine-bridge configurations defined in Figure 5 and Supplementary Figure S8. These labels describe ENV structural architecture rather than formal phylogenetic or taxonomic groups. Several errantivirus HSV/gB-type ENV groups showed comparable or lower levels of primary-sequence identity while retaining lineage-specific cysteine-bridge architectures, including Cnidaria elements with the C1c configuration (18.1–32.2%), Ctenophora elements with the C1c configuration (21.5–29.5%), Bryozoa elements with the C2b configuration (median 28.1%), Branchiopoda elements with the C2c configuration (median 27.1%), Tardigrada elements with the C2c configuration (median 26.5%) and Nematoda elements with the C1a/d configuration (median 19.9%). Thus, conserved cysteine-bridge architecture in errantivirus HSV/gB-type ENV groups cannot be explained simply by recent duplication or high overall sequence identity. Rather, it supports long-term structural constraint on lineage-specific ENV architectures despite extensive primary-sequence divergence. These pairwise identities are used here as descriptive measures of sequence divergence, not as calibrated molecular-clock estimates.

The configuration of cysteine bridges appears to be specific to phyla, though this does not seem to be reflected by their positions in the tree based on POL extended RT/connection tree. For example, Bryozoa elements (Tree positions A3 and A9 in fig S4B) all carry configuration ‘C2b’, despite being found in different clades on the tree, even though they each cluster themselves separately from other elements (Fig. S4B, 5D and S8). Similarly, errantiviruses carrying HSV/gB-type *env* found in insect phyla mostly have the same configuration regardless of their position in the POL extended RT/connection tree. For example, insect elements within the “ancient errantivirus” group, and insect elements within the “insect errantivirus” group, all have the same cysteine bridge configuration in HSV/gB (Fig 5D and S8). There are, however, cases where more than one type of configuration can be found within the same phyla, namely Nematoda and Platyhelminthes (Fig 5D). Though, in those cases, they appear to continue to separate based on POL extended RT/connection tree (Fig. S8). For example, Nematoda elements with configuration ‘C2a’ and ‘C2c’ are found in a different clade to those with ‘C1a’ and ‘C1d’ (Fig 5D). Although, it should be noted that elements with ‘C1a’ in Nematoda are found in two different clades (Fig 5D).

As mentioned previously, the clades that form on POL extended RT/connection mostly separate elements that carry HSV/gB-type ENV from F-type ENV within discrete clades (Fig. S4B). This, with the fact that the same substructures can be found in different orders, for example in Nematoda, Bryozoa and Platyhelminthes, suggests that their origin is likely ancient. The overall specificity of certain residues to host orders further suggests that recombination events would have only occurred between elements within the same animal order. Furthermore, the overall separation of distinct cysteine bridge configurations of HSV/gB-type ENV, that are represented in the POL extended RT/connection tree within ancient phyla, suggests that those recombination events occurred anciently. While insect elements are separated on the tree, ERVs with HSV/gB-type ENV tend to cluster separately from those with F-type ENV. For example, in the “insect errantivirus” group, the clade that almost exclusively carries HSV/gB-type ENV sits sister to another clade that only carries F-type ENV (Fig 3B). Not only do they separate from each other, but HSV/gB-type ENV can be found in a variety of superfamilies within Hymenoptera, which suggests their ancient origin (Fig 3B). Some species, such as a hymenopteran *Lasioglossum lativentre*, carry elements with HSV/gB-type, as well as those with F-type ENV (Fig 3B and Table S1). Despite being found in the genome of the same species, the *pol* of errantiviruses with F– and HSV/gB-type ENV do not stay together, further supporting their more ancient origin (Fig 3B). It should be noted that some clades, such as ones in tree position A2 within “ancient errantiviruses”, have elements that carry F– and HSV/gB-type ENV sitting together within the same discrete clade, suggestive of more recent recombination (Fig. S4B).

### The RNase H-containing bridge region of *pol* separates “insect errantiviruses” from “ancient errantiviruses”

To further characterise the errantiviruses identified in this study, we next asked whether Pol domain architecture differed between the major phylogenetic groups. We focused on the RNase H-containing bridge region, which we define here as the region spanning the canonical RNase H domain and the C-terminal region between RNase H and Integrase. This region is distinct from the extended RT/connection region used for the Pol phylogenetic analyses above. The connection subdomain included in the extended RT/connection alignment corresponds to a partial RNase H-like fold, whereas the canonical RNase H domain analysed here contains the complete β1–β2–β3–α–β4–α–β5–α architecture shown in Figure 6A. Because RNase H-related domains have changed during retroelement evolution, including the duplicated and partially degraded RNase H-like tether/connection domain described in vertebrate retroviruses (Malik and Eickbush 2001), the bridge region provides an additional structural feature for comparing errantivirus lineages. We therefore characterised the bridge region of representative elements using AlphaFold 2, focusing on the organisation of the canonical RNase H domain and the downstream RNase H-like or mini-domain structures.

We found various structures within the bridge region, mostly consisting of core β-sheets resembling the 3D fold arrangement of RNase H domains but we also found other uncommon substructures (Fig 6A). Almost all errantiviruses that sit within the “insect errantivirus” group have duplicated the RNase H domain similarly to vertebrate retroviruses, with the C terminal copy presenting itself as a “pseudo” RNase H domain (Fig 6A-B). This “pseudo” RNase H domain retains the barrel-like structure made of five core β-sheets, but the order of the second and the third β-sheets is flipped (Fig 6A).

On the other hand, “ancient errantiviruses” carry an RNase H domain, and a C-terminal “mini” domain. This consists of three small core β-sheets and α-helix residues at its N terminus. This “mini” domain is also carried by insect elements in this group, further suggesting that those insect elements are equally ancient as the other errantiviruses in this group. Furthermore, this “mini” domain is also found in the bridge region of yeast *S. pombe tf1 pol* (Fig 6B-C), indicating it as a feature of highly ancient transposons. Thus, Figure 6 links the POL extended RT/connection phylogenetic separation of insect and ancient errantiviruses with a structural difference in the bridge region of Pol.

The other structures (“Mini” domains 2, 3 and “Pseudo” domain 2) present themselves less frequently as outliers, and are restricted to specific clades, or even solitary elements (highlighted in yellow and dark blue in Fig 6B). One notable outlier is an insect element within the “ancient errantivirus” group that carries a variation of a “pseudo” RNase H domain (position 7, noted as “pseudo” domain 2 in Fig 6). This same domain can be found in a phylogenetic group in the “insect errantivirus” group (position 3 in Fig 6B). Furthermore, some elements in this “insect errantivirus” group carry a “mini” domain as well (position 3 in Fig 6B). Another is that some elements within the “insect errantivirus” group only carry a solitary RNase H domain (positions 4 and 5 in Fig 6B). It should be noted that these outliers are mostly specific to small phylogenetic groups/clades, and likely also arose anciently.

### Intact errantiviruses are sourced from various lineages of *Ty3/gypsy* retrotransposons

To further investigate the origins of *env*-containing errantiviruses, we assembled a phylogenetic tree from the extended RT/connection region of representative errantiviruses and included non-*env-*containing retrotransposons that fall under the broader umbrella of “*Ty3/gypsy*” elements. This includes transposons such as *MAG* and *Tor*, and Chromoviruses such as *Maggy*, which are annotated in the GypsyDB (https://gydb.org). We expected that *env*-containing errantiviruses would form a monophyletic clade in the tree if they arose from a specific non-*env-*containing *Ty3/gypsy* element.

We firstly find that most non-*env*-containing *Ty3/gypsy* elements sit in the ‘ancient errantivirus’ clade, with the exception of the *412/MDG1* group – specifically the Drosophila *gypsy* elements *BLOOD* and *Gypsy-28_Dan* – which cluster with Drosophila and other insect errantiviruses, consistent with a previous study (Bargues and Lerat 2017) (Fig 7). Additionally, we find that the non-*env-*containing elements rarely regularly form discrete clades without *env-*containing errantiviruses. For example, *Tor4b* is found sister to Tunicata elements carrying F-type *env*, and *Cer1* was found sister to a Platyhelminthes errantivirus carrying HSV/gB-type *env*. Interestingly, Chromoviridae, which are found in Fungi and Plants and are representative of highly ancient, widespread *Ty3/gypsy* elements (Gorinsek et al. 2004; Kordis 2005), are found to cluster within the tree, nested with “ancient” errantivirus clades (Fig 7). They sit within a clade that includes Tunicata errantivirus carrying HSV/gB-type *env*.

**Figure 7.**
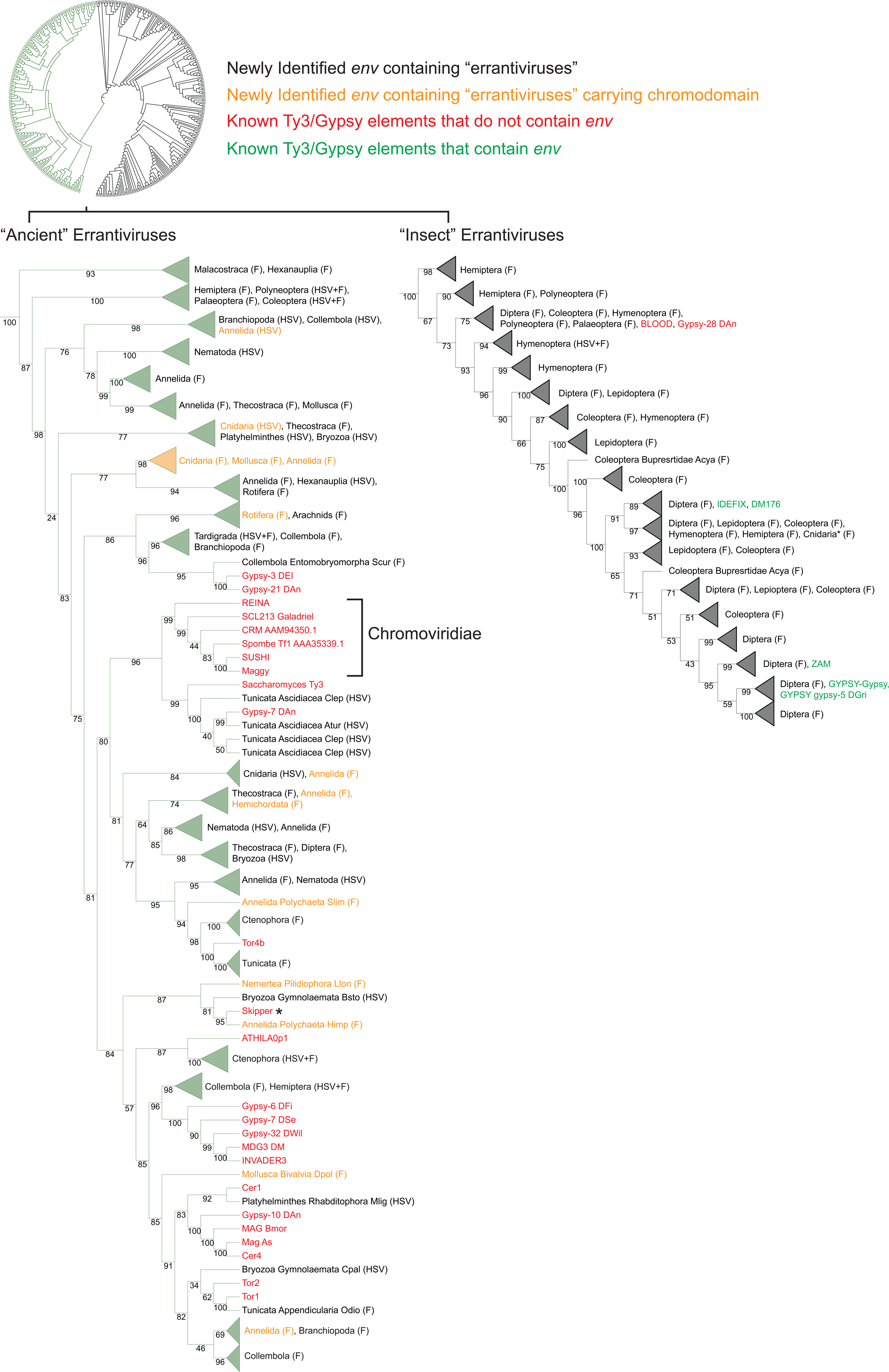
– Trees of known *Ty3/gypsy* elements and newly identified Errantiviruses. Midpoint-rooted maximum-likelihood tree of the POL extended RT/connection region from representative *env*-containing errantiviruses and selected non-*env*-containing *Ty3/gypsy* retrotransposons. Representative non-*env*-containing *Ty3/gypsy* elements from animals, fungi and plants are included to place the newly identified errantiviruses within the broader diversity of *Ty3/gypsy* retrotransposons. Host Order and *env* type given for each clade, as well as solitary elements on the tree are shown. Errantiviruses containing chromodomains are marked in yellow. The “ancient errantivirus” group is coloured in green and “insect errantiviruses” in black. Known *Ty3/gypsy* elements includes Gypsy Group 1 OSIRIS (Gypsy-21_DAn), Gypsy Group 1 OSVALDO (Dypsy-3_DEl), Gypsy Group 2 BICA (Gypsy-6_DFi), Gypsy Group 2 BLASTOPIA (Gypsy-32_DWil), Gypsy Group 2 MDG3 (INVADER3, MDG3_DM), Gypsy Group 2 MICROPIA (Gypsy-7_DSe), Gypsy Group 2 SACCO (Gypsy-10_DAn), Gypsy Group 3 412/MDG1 (BLOOD, Gypsy-28_Dan), Gypsy Group 3 CHIMPO (Gypsy-7_DAn), Gypsy Group 3 17.6 (DM176, IDEFIX, ZAM), Gypsy Group 3 GYPSY (Gypsy-5_Dgri, Gypsy-Gypsy), Mag (Mag_As, MAG_Bmor), Cer (Cer1, Cer4), Tor (Tor1, Tor2, Tor4b), Athila/Tat (ATHILA0p1), and Chromoviridae (ATHILA0p1, CRM_AAM94350.1, REINA, Saccharomyces_Ty3, SCL213_Galadriel, Skipper*, Spombe_Tf1_AAA35339.1, SUSHI).

The annotated *Ty3/gypsy* elements present themselves in the tree, spread between and within different “ancient” errantiviruses from different hosts, with little pattern to their spread regarding host or *env* type. Notably, a clade of errantiviruses from Cnidaria, Mollusca and Annelida hosts all carry a chromodomain, a chromatin-binding domain found C-terminal to Integrase in chromoviruses and some related *Ty3/gypsy* elements. This clade sits basally to known chromoviruses, but sister to non-chromodomain-containing errantiviruses (Fig 7, Fig. S9). We also observe outliers, where only one or two errantiviruses carry chromodomains within the whole clade. For example, Annelida_Polychaeta_Dgyr_Errantivirus_1 contains a chromodomain, but is located in a clade with non-chromodomain-containing errantiviruses (Fig. S9). These observations rule out a single origin of *env-*containing errantiviruses and instead suggest that multiple lineages of ancient *Ty3/gypsy* elements contributed to the current distribution and diversity of errantiviruses.

## Discussion

We found copies of *env-*containing *Ty3/gypsy* retrotransposons (errantiviruses) in the genomes of nearly every major metazoan order. They are intact, carrying uninterrupted ORFs of all three retroviral proteins and are flanked by LTRs. Additionally, most errantiviruses identified in this study have multiple copies in their respective genomes, consistent with recent or ongoing genomic expansion. When we further characterized these elements, we found that they carry different ENV proteins; ENV resembling glycoprotein F from paramyxoviruses, and ENV resembling glycoprotein B from herpesviruses. Despite great divergences in the primary sequence, the structural domains of these proteins are highly conserved. Throughout this manuscript, we have used “errantivirus” as an operational term for *env*-containing *Ty3/gypsy* retrotransposons, including elements outside Drosophila. This usage should not be interpreted as a formal taxonomic proposal, nor as evidence that all such elements share the same amplification mechanism as Drosophila *gypsy*. Rather, it reflects their shared genomic organisation, *Ty3/gypsy* ancestry and possession of *env*-like fusogen genes.

We found that these errantiviruses are highly ancient, being found in animals such as cnidarians, ctenophores and tunicates – lineages that diverged early in metazoan evolution. We identified an “ancient errantivirus” group, which sits separately to elements found in insects (Fig 2). These ancient elements are also structurally distinct, with the bridge region between RT and IN of *pol* resembling that of yeast *Ty3* retrotransposons, rather than that of insects. Additionally, *pol* genes of non-*env*-containing *Ty3/gypsy* elements from diverse eukaryotic taxa form clades with the “ancient errantivirus” group (Fig 7). Although these patterns are consistent with their ancient nature, the specific origins of *env-*containing errantiviruses in metazoans remain elusive. Errantiviruses from the same or closely related phyla tend to cluster together in the POL extended RT/connection phylogenetic tree (Fig. S4). However, the pattern becomes convoluted when highly ancient errantiviruses are compared, as elements from distantly related phyla often cluster together without representation from intermediate animal lineages (Fig. S4 and Fig 7). Many of these clades tend to have longer branches, suggesting that they are distantly related and diverged a long time ago. For example, within tree position A10 (Fig. S4), a clade consisting of elements from Hexanauplia and Annelida are separated by longer branch lengths. The reason for these types of clades, and as to why we do not see elements in species between those distantly related orders, is likely due to the frequent loss of these retrotransposons from the genome. In such cases, distantly related species may retain intact or recently expanded copies that were vertically inherited and diverged with the host species, while copies in other species may have become inactive and degraded over time.

The errantiviruses we identified form multiple clades with previously characterized non-*env-*carrying *Ty3/gypsy* elements in the *pol* tree (Fig 7). Additionally, some errantiviruses carry chromodomains, but do not always cluster together with other chromodomain carrying retrotransposons. This pattern could be explained by different lineages of *Ty3/gypsy* frequently acquiring *env* from other errantiviruses. This would explain annotated *Ty3/gypsy* elements forming clades with both HSV/gB– and F-type *env*-containing errantiviruses. Another explanation could be that widespread horizontal transfer is occurring between distantly related species. As the species diverge from each other, different types of *env* could be acquired, lost and regained multiple times. However, neither of these explanations account for the congruence seen between *pol* and the host, at least at the level of animal classes and orders (Fig. S4), nor the structural divergence of HSV/gB-type ENV (Fig 5 and S8). The spread of errantiviruses is indicative of multiple lineages of retrotransposon contributing to the current picture of errantiviruses, but the congruence supports their ancient nature. One plausible explanation is that different lineages of *Ty3/gypsy* within the last common ancestor acquired *env*, which were inherited and diverged with the host species, infrequently recombining with other elements within the same/similar hosts. The non-canonical GAG-ENV-POL arrangement further supports this view. These elements are found in several non-insect animal groups and do not form a single monophyletic clade, indicating that this arrangement is not simply a recent derivative of one lineage (Fig. S4 and S8). Instead, ORF rearrangement appears to be another feature that has diversified among long-standing HSV/gB-type errantivirus lineages.

Our interpretation does not exclude lineage-specific loss or rare horizontal transfer. Indeed, extensive loss is expected for ancient retrotransposon lineages, because inactive copies will progressively degrade and disappear from detectable sequence space. Similarly, occasional horizontal transfer between closely related hosts may contribute to local discordance. To examine whether host-taxonomic structure is nevertheless retained at local evolutionary scales, we performed targeted host-label permutation analyses of selected Annelida and Lepidoptera POL RT subclades (Fig. S5). The Annelida subclade showed strong element–host concordance at both host-family and host-species levels. The Lepidoptera subclade showed a more mixed pattern, with several highly supported clades enriched for related host groups at the superfamily or broader taxonomic level, but also a substantial number of mixed clades. Thus, lineage-specific loss and rare horizontal transfer are compatible with our data, but frequent broad horizontal transfer between distantly related animal groups is unlikely to explain the overall host-taxonomic structure.

### Acquisition of ENV proteins by errantiviruses

It was originally speculated that errantiviruses acquired *env* from baculoviruses in insects, some of which carry an F-type ENV protein Ld130, potentially via recombination events (Malik et al. 2000; Rohrmann and Karplus 2001). While *Ty3/gypsy* elements may have obtained the *env* gene via recombination, it is unlikely that baculoviruses are the source for the *env* gene found in errantiviruses, as they are far more widespread than insects. The congruence of phylogeny between *env* and *pol* indicates that they have likely co-evolved (Fig 4). This, along with the presence of errantiviruses in ancient host orders such as Cnidaria, Ctenophora and Tunicata, suggests that *env*-carrying *Ty3/gypsy* elements are highly ancient transposons, rather than the result of a recent recombination event with an exogenous virus. HSV/gB-type *env* in errantiviruses appears to be equally ancient, as it is similarly found in ancient animal orders (Fig 1). The arrangement of cysteine bridges that stabilize the structure of the ectodomain, are consistent between elements within the same order, even though *pol* presents itself separately on the tree (Fig 5 and S8). This implies that the *env* has diverged with the host but likely undergoes recombination events with other elements within the same species. Though, *pol* genes with HSV/gB-type *env* mostly cluster within their own clades and are not mixed with *pol* with F-type *env*, indicating that recombination events are not happening frequently enough to randomise their spread, and likely occurred anciently. The origins and acquisition, therefore, of either type of *env*, likely occurred in the last common ancestor of metazoans.

### The survival mechanism of errantiviruses in the genome

In Drosophila, errantiviruses occupy the somatic cell niche in the ovaries that surround the germline, and the expression of *env* allows virus-like particles to infect the germline to create new copies (Song et al. 1994). As an adaptation, Drosophila species have duplicated the germline transposon defence mechanism to target errantiviruses in the somatic cells of the gonad (Brennecke et al. 2007). We recently showed that errantiviruses are targeted by piRNAs in ovarian somatic cells across multiple insect orders, including mosquitoes, bees and crickets (Belot et al. 2020). It is therefore possible that this is more universal, and that *env*-containing retrotransposons occupy the cell niches surrounding the germline, and that expression of *env* allows them to continue to infect the germline. For example, murine IAP retrotransposons that carry *env* genes also remain infectious, like Drosophila errantiviruses (Ribet et al. 2008). However, these additional ORFs and genes in endogenous retroviral elements are constantly subject to evolutionary pressure from their hosts (Wang and Han 2020; Wang et al. 2020). It is therefore significant that the *env* gene is as highly conserved as we have observed, and that it does not seem to have been subject to frequent loss and acquisition. It is also important to note the mechanisms of transmission via the action of this *env* gene remain elusive for most of the elements we identified, except for those in Drosophila species. However, the structures formed by *env*, for both HSV/gB and glycoprotein F, form the same functional structures as those found in exogenous viruses, required for cell entry (Heldwein et al. 2006; Connolly et al. 2011; McLellan et al. 2013). While entry of HSV also requires three other glycoproteins, gD, gH and gL, gB is specifically required for membrane fusion (Johnson et al. 2011; Jambunathan et al. 2021). The mechanism of cell entry, therefore, is unknown for these errantiviruses that carry gB, and necessitates further investigation regarding their expression. Though, given their highly structured configuration, it is likely that they retain membrane fusion capabilities. Additionally, given the widespread nature of these *env*-containing elements, it may also be possible that somatic transposon defence mechanisms against them are more common.

In summation, the *env* gene appears to have been anciently acquired by *Ty3/gypsy* elements, and has been retained by those elements, diverging as their host species evolved. Today, we see a snapshot of errantiviruses widespread throughout metazoan genomes, continuing to create new insertions to survive long term.

## Materials and methods

### Genome assemblies used in the study

Reference genome sequences were retrieved from NCBI and grouped into the following insect orders and other metazoan phyla: Diptera, Lepidoptera, Coleoptera, Hymenoptera, Hemiptera, Polyneoptera, and Palaeoptera for insects; Arachnida, Branchiopoda, Chilopoda, Collembola, Diplopoda, Hexanauplia, Malacostraca, Merostomata, Ostracoda, and Thecostraca for Arthropoda; Acanthocephala, Annelida, Brachiopoda, Bryozoa, Dicyemida, Mollusca, Nemertea, Onychophora, Orthonectida, Phoronida, Platyhelminthes, Priapulida, Rotifera, and Tardigrada for Protostomia; Cephalochordata, Echinodermata, Hemichordata, and Tunicata for Deuterostomia; Cnidaria, Ctenophora, and Porifera. We obtained lists of reference genomes in each category in May 2023 using the Entrez Direct command line tool. The list of the genome assemblies used can be found in Supplementary Table S6.

### Identification of intact genomic copies of *Ty3/gypsy* errantiviruses

We ran a total of five iterations of tBLASTn (ncbi-blast-2.9.0) to search for the genomic copies of *Ty3/gypsy* errantiviruses, firstly using the baits of known elements, and subsequently using the elements that were identified after each iteration. We chose new baits that were not represented in the phylogenetic tree of the POL (see below for the detail) from the previous iteration and repeated the process until we no longer found new elements. We used GAG, POL and ENV protein sequences of the Dipteran *gypsy* errantiviruses from the RepBase (https://www.girinst.org/repbase/) for the first tBLAST search. In this search pipeline, “ENV” is used as a generic term for candidate envelope-like proteins. Candidate ENV ORFs were classified after identification as either F-type ENV or HSV/gB-type ENV based on sequence similarity, domain prediction and structural features. The initial ENV baits were derived from Dipteran *gypsy* errantiviruses, whereas subsequent iterations also used ENV proteins from newly identified representative elements, allowing both F-type and HSV/gB-type ENV-containing elements to be recovered. The bait sequences for all tBLASTn iterations can be found at https://github.com/RippeiHayashi/errantiviruses_2025/tree/main. We ran tBLASTn with default parameters (-evalue 10). We applied different criteria to identify ERVs from species outside insects as we did not have prior knowledge of whether or what kind of ERVs are found in those species. We briefly summarise our approach below.

Genomic regions that showed homology to the POL baits with a bit score of greater than 50 or to the GAG and ENV baits with a bit score of greater than 30 were individually pooled. Genomic regions that had hits from multiple baits were merged for their coordinates using bedtools/2.28.0. Regions that were larger than 1500 nucleotides for POL and 300 nucleotides for GAG or ENV were collected. Following, for insect genomes we looked for regions that carry GAG, POL, ENV sequences that are next to each other in this order allowing up to 500 nucleotides of gaps between them as well as excluding regions of smaller than 4000 nucleotides or larger than 10000 nucleotides. For the other genomes we allowed up to 1000 nucleotides of gaps between POL and ENV and collected regions that had both ORFs that are not smaller than 3000 nucleotides or larger than 10000 nucleotides.

We subsequently checked whether the regions contain uninterrupted ORFs by making translated sequences using transeq from EMBOSS-6.6.0 and excluding ORFs with more than five stop codons. We clustered all the identified GAG, POL and ENV from each genome by Clustal Omega (https://www.ebi.ac.uk/Tools/msa/clustalo/) and identified genomic regions that harbour all three ORFs that are multiplied in the genome. We also searched for ORFs regardless of the copy number when there were no multiplied elements in the genome. When genomic copies were abundantly found in multiple closely related species, we only annotated the elements from representative genomes. Genomes that were annotated and the number of predicted ORFs per genome are summarised in Supplementary Table S6.

These length, gap and stop-codon thresholds were used as permissive prefilters to recover candidate elements for downstream curation, rather than as the sole definition of intact errantiviruses. We chose inclusive thresholds at this stage to avoid excluding divergent elements, elements with non-canonical ORF arrangements, or elements with small annotation errors. Final inclusion required additional evidence, including *Ty3/gypsy* POL classification, recovery of full-length RT and Integrase domains within continuous ORFs, HHpred-supported domain annotation of GAG, POL and ENV, and removal of elements with large domain truncations.

The above process yielded abundant putative intact copies of ERVs from insect genomes whereas we took additional steps for the other genomes. For Arthropoda genomes outside insects as well as other Protostomia genomes, we extended the genomic regions of predicted GAG-POL to downstream by 3000 nucleotides and manually annotated ENV ORFs using Benchling (https://www.benchling.com/). The identity of GAG ORFs were further confirmed by HHpred (https://toolkit.tuebingen.mpg.de/tools/hhpred). For Cnidaria, Ctenophora and Deuterostomia genomes outside vertebrates we used transeq to annotate putative GAG and ENV ORFs from 5000 nucleotides downstream and 3000 nucleotides upstream to the predicted POL ORFs, respectively.

We took confident 945 POL ORFs from the first tBLASTn search and made a multiple sequence alignment of the conserved regions of the Integrase domain using mafft-7.505 with the option ‘--auto’, and inferred the phylogenetic relationships using iqtree-1.6.12 with the option ‘-m rtREV+R4 –bb 1000’ (see more details in the section below). We took 152 representative POL from the phylogenetic tree and used the POL and the corresponding GAG and ENV ORFs from the same genomic regions for a second tBLASTn search. We repeated the above steps after combining the results of the two tBLASTn searches. We did not find any ERVs in Nematoda genomes that have GAG, POL and ENV in this order while the previous study reported non-canonical Nematoda ERVs that have flipped the order of ENV and POL (Sacco et al. 2022). Thus, we searched for regions that have ENV and POL in this order allowing gaps of up to 2000 nucleotides between them. GAG was subsequently predicted from 5000 nucleotides upstream to the predicted ENV-POL. We extended the search for potential GAG-ENV-POL insertions to other non-insect genomes and found similar elements in Bryozoa and Platyhelminthes. We also found ENV-GAG-POL elements in Arachnida genomes (*Medioppia subpectinata* and *Oppia nitens*). These elements were recovered through the search of ENV-POL elements because GAG ORF was short. We did not extend our search to find other arrangements of ORFs.

Additionally, we found that the glycoprotein gene present in mosquito *BEL/pao* retrotransposons (Dezordi et al. 2020) are also found next to *gypsy* POL genes. The glycoprotein gene resembles the *envelope* genes from Chuviridae and is structurally similar to glycoprotein B from Herpesviruses. We identified ten mosquito *BEL/pao* retrotransposons that carry this glycoprotein from RepBase (BEL-199_AA, BEL-279_AA, BEL-652_AA, BEL-801_AA, BEL-804_AA, BEL-879_AA, BEL-51_AnFu, BEL-56_AnFu, BEL-81_AnFu, and BEL-71_CQ) and used them as baits for a tBLASTn search against Arthropoda genomes. The tBLASTn search identified 114 *gypsy* elements with homologous glycoprotein genes from Coleoptera, Hymenoptera, Polyneoptera, and Hexanauplia species. Combined together, the second iteration of tBLASTn search identified 451 new elements. The phylogenetic analysis of POL from the elements identified in the first and second tBLASTn runs revealed clades that only contained new elements from the second tBLASTn run, suggesting that the first two tBLASTn runs were not comprehensive. Therefore, we took 34 additional baits to cover all the major clades in the phylogenetic tree for a third tBLASTn search, which identified 61 new elements. We repeated tBLASTn runs five times in total; the fourth and fifth runs used 17 and 7 new baits and identified 49 and 6 new elements, respectively. We stopped the tBLASTn search after five iterations because the phylogenetic analysis of POL did not reveal new clades.

Following, we further curated newly identified ERVs. Firstly, we used hmmer 3.3.1 (http://hmmer.org/) to make sure that the ERVs that we identified are classified as *Ty3/gypsy*. We compared POL ORFs from the newly identified ERVs against the POL ORFs of known retroelements that are available from the Gypsy Database 2.0 (https://gydb.org/index.php/Main_Page). The Gypsy Database 2.0 collects hmm profiles of *Ty1/Copia*, *Bel/Pao*, *Ty3/gypsy*, and Retrovirus POL proteins. The hmmer search classified all the newly identified ERVs into *Ty3/gypsy*. Secondly, we ran HHpred from hh-suite 3.3.0 (Steinegger et al. 2019) or the online tool of HHpred (https://toolkit.tuebingen.mpg.de/tools/hhpred) to predict GAG, ENV, and POL ORFs using the default search parameters. For the command line HHpred, we first made multiple sequence alignments using hhblits and the Uniclust database UniRef30/2022_02 (https://uniclust.mmseqs.com/). This was followed by hhsearch to predict domain structures using the collection of pdb70 from March 2022 (https://wwwuser.gwdg.de/∼compbiol/data/hhsuite/databases/hhsuite_dbs/). We used default settings to run hhblits and hhsearch. ENV and POL ORFs with large truncations were removed after the domain structure prediction. ERVs that predicted all GAG, ENV and POL ORFs were taken for further analyses.

We additionally ran BLASTn using the genomic regions that cover all three ORFs as baits to interrogate the copy number of ERVs in the genome where they are found. The results are summarised in Supplementary Table S1 where we counted the number of BLASTn hits with a more than 98% identity in more than 98% of their length to the bait sequences. A majority of newly identified ERVs (1386 out of 1512 elements) carry ORFs that are found at least twice in the genome. These copy-number estimates were used as evidence consistent with recent or ongoing genomic expansion, but were not taken as direct evidence of current transposition activity.

For selected Porifera and Echinodermata genomes in which no intact GAG-POL-ENV elements were detected, we additionally searched for full-length GAG-POL *Ty3/gypsy* elements to assess whether intact retrotransposons could be recovered from these assemblies. Two representative genomes were analysed from each phylum. Candidate regions containing GAG and POL in the expected order were identified and curated using the same general criteria applied to GAG-POL-ENV elements, except that no downstream ENV ORF was required. Candidate GAG-POL elements were then assessed for copy number, putative LTRs and tRNA primer-binding sequences as described above.

### Identification of putative long terminal repeats and transfer RNA (tRNA) primer binding sequences

Long terminal repeats (LTRs) of ERVs were identified as follows. Firstly, we fetched the genomic coordinates of ERV insertions and extended 3000 nucleotides at either ends from the Start codon of GAG and the Stop codon of ENV ORFs (POL was taken for the GAG-ENV-POL ERVs) for all species categories except for Cnidaria, Ctenophora and Deuterostomia where we extended 5000 nucleotides at either ends from the Start and Stop codons of POL. We retrieved the genomic sequence of the extended ERVs and ran BLASTn against the same sequence to identify repeated sequences. We considered repeated regions of greater than 99% identity and the tBLASTn bit score between 200 and 5000 as putative LTRs. We excluded LTRs whose distance between them is shorter than 5000 nucleotides or longer than 14000 nucleotides. Following, we searched for putative tRNA primer binding sequences (PBS). tRNA sequences were retrieved from genomic annotations from representative genomes in respective species categories except for Ctenophora where we took tRNA sequences from Protostomia, Cnidaria and Deuterostomia because there were no annotated genomes available at the time of the study. Representative genomes that we used to retrieve tRNA sequences are: GCF_000001215.4 and GCF_013141755.1 for Diptera, GCF_014905235.1 and GCF_905147365.1 for Lepidoptera, GCF_000002335.3 and GCF_917563875.1 for Coleoptera, GCF_003254395.2 and GCF_016802725.1 for Hymenoptera, GCF_014356525.2 and GCF_020184175.1 for Hemiptera, GCF_002891405.2 and GCF_021461395.2 for Polyneoptera, GCF_921293095.1 for Palaeoptera, GCF_000671375.1, GCF_020631705.1, GCF_002217175.1, GCF_016086655.3, GCF_024679095.1, GCF_019059575.1 for Arthropoda, GCF_000326865.2, GCF_020536995.1, GCF_000002985.6, GCF_000715545.1 for Protostomia, GCF_000222465.1 and GCF_022113875.1 for Cnidaria, GCF_013122585.1, GCF_000003605.2, GCF_000003815.2, and GCF_000002235.5 for Deuterostomia. The minimum requirements of the priming events are incompletely understood, but the length of PBS for retrotransposons is generally between 8 and 18 nucleotides and the PBS is positioned near the 3’ end of the 5’ LTR (Wilhelm and Wilhelm 2001). We chose the 12 nucleotides at the 3’ end of tRNAs as putative primer sequences and searched for PBS in the 100 nucleotide-window centred at the 3’ end of the 5’ LTR using bowtie-1.2.3 allowing up to two mismatches. We added “CCA” to the 3’ end of the tRNA sequence when it is not present in the genomic sequences. We successfully identified putative PBSs in 1151 out of 1512 ERVs that we identified in this study. We grouped ERVs based on the type of tRNA used for the PBS and summarised the results in Figures S2 and S4 as well as Supplementary Table S1.

### Phylogenetic analysis of POL

We obtained multiple sequence alignments of the POL extended RT/connection region and Integrase domains as follows. For clarity, we distinguish between the extended RT/connection region used for phylogenetic analysis and the RNase H-containing bridge region analysed structurally in Figure 6. The extended RT/connection region comprises the RT polymerase core, formed by the fingers, palm and thumb subdomains, together with the downstream connection subdomain. Although the connection subdomain is included as part of retroviral RT in structural studies, it corresponds to a partial RNase H-like fold and has been interpreted in evolutionary studies as a degenerated RNase H-like tether domain (Malik and Eickbush 2001). In contrast, the bridge region analysed in Figure 6 is defined here as the region spanning the canonical RNase H domain and the C-terminal region between RNase H and Integrase. In the structural scheme used in Figure 6A, an intact RNase H domain consists of β1–β2–β3–α–β4–α–β5–α, whereas the connection/tether region included in the extended RT alignment corresponds only to the partial RNase H-like fold β1–β2–β3–α–β4–α.

For the POL extended RT/connection region, we first took the HHpred alignments of POL against the RT structures of Moloney Murine Leukaemia Virus (Protein Data bank entry 4MH8_A) and Human Immunodeficiency Virus type 1 (Protein Data bank entry 4G1Q_B), spanning regions that correspond to their fingers, palm, thumb and connection subdomains (Das and Georgiadis 2004), namely the amino acid positions of 28 to 426 and 7 to 405, respectively. Thus, the region used for this phylogenetic analysis is broader than the RT polymerase core alone, but does not correspond to the complete canonical RNase H domain analysed structurally in Figure 6. This region was used because it is conserved and alignable across the errantivirus POL proteins analysed here. However, this process caused small truncations at either ends of the domain because of poor alignments at the individual substructures. To overcome this issue and to facilitate sequence alignments, we extended the N– and C-termini by 10 and 15 aa, respectively, after merging the two predicted regions of RT domains. We then ran mafft-7.505 to generate a multiple sequence alignment, from which we manually trimmed the either ends to have intact fingers, palm, thumb and connection subdomains for all POL ORFs. Similarly, we took HHpred alignments of POL against the Integrase domains of Human Spumaretrovirus (Protein Data bank entry 3OYM_A) and Simian T-lymphotropic Virus type 1 (Protein Data bank entry 7OUH_B), spanning regions that correspond to their N-terminal domain (NTD) and catalytic core domain (CCD) (Engelman and Cherepanov 2014), namely the amino acid positions of 54 to 273 and 8 to 197, respectively. We used the predicted Integrase domains with 15 and 30 aa extensions at the N– and C-termini, respectively, for generating a multiple sequence alignment before manually trimming the ends. We inspected the multiple sequence alignments using Mview (https://www.ebi.ac.uk/Tools/msa/mview/) and removed ERVs that contained truncated RT or Integrase domains from the analyses. We included ORFs from 5 Drosophila errantiviruses (DM176, IDEFIX, ZAM, GYPSY, and Gypsy-5_DGri) in the multiple sequence alignments.

We ran iqtree-1.6.12 with the option ‘-m rtREV+R4 –bb 1000’ to generate the phylogenetic trees of all POL Integrase domains and extended RT/connection regions, and those from ERVs found in vertebrate and invertebrate genomes, respectively. We then selected representative elements from clades, taking into consideration the relative distances between nodes in the same clade. For example, if the distance between two nodes is less than 0.1, meaning that there is little – no variations between their sequences, only one node in that position is taken. Additionally, we made sure that the representatives captured the diversity of *env* found within the clade. For example, if a clade contains both F-type and HSV/gB-type elements, at least one for both is taken, regardless of distance. Furthermore, the phylogeny of the hosts was represented as fully as possible. For example, if a clade has elements from Diptera and Coleoptera, at least one from both is taken regardless of distance. Once representatives were collected, the tree was then accordingly pruned. It should be noted that the final pruned tree of representatives was only used as a figure aid for clarity (Fig 1-4), and all analyses were done on the full Integrase and extended RT/connection phylogenetic trees. We used a similar approach as detailed to select representatives for structural analysis of *env* ectodomains, as well as the *pol* bridge region (Table S4). If any outliers were found, remaining elements from the same clade were also analysed, to check whether structural inconsistencies were consistent across the clade.

For congruence plots in figures 4 and S2, we grouped elements based on which node they sit in the tree, ensuring that the node is well supported by bootstrap values greater than 95. We used the networkD3 package in R to generate Sankey plots to visualise the representation of elements from one tree in another. The node definitions are summarized in Table S5.

Further, we took 86 representatives of the F-type ENV based on their positions in the corresponding POL phylogenetic tree for the analysis. We identified the signal peptides and transmembrane regions by DeepTMHMM (https://dtu.biolib.com/DeepTMHMM) and obtained the ectodomains that are in between them. We subsequently ran mafft-7.505 to generate a multiple sequence alignment. To compare errantivirus F-type ENV proteins with homologous fusogens outside *Ty3/gypsy* retrotransposons, we generated an additional F-type ENV ectodomain alignment that included representative external sequences. These included baculovirus F proteins, paramyxovirus and pneumovirus F proteins, and the BEL/Pao-associated *Drosophila melanogaster* Roo F-like protein.

### Host-taxonomic concordance analyses

For targeted host-taxonomic concordance analyses, we selected representative Lepidoptera and Annelida POL extended RT/connection subclades from tree positions I1 and A6, respectively, and scored internal nodes with bootstrap support ≥95 and 3–10 descendant tips. Host-taxonomic labels were randomly permuted across the fixed POL extended RT/connection topology 10,000 times while preserving the observed label counts, and the same supported nodes were rescored after each permutation. In the Lepidoptera analysis, family labels were permuted and superfamily assignments were derived from the permuted labels. In the Annelida analysis, host-species labels were permuted and host-family assignments were derived from the permuted labels.

### Structure prediction of ENV proteins, the RNase H and chromodomains using Alphafold 2

We took representatives of the F-type and HSV/gB-type ENV proteins and the RNase H-containing bridge region, defined here as the canonical RNase H domain and the C-terminal region between RNase H and Integrase, using branch definitions from the phylogenetic trees of POL extended RT/connection region. We also took the C-terminal half of POL, including the Integrase domain, from representative errantiviruses, investigated in Figure 7. We predicted their structures when chromodomains were detected by hhpred (see above). We used the colabfold wrap v1.4.0 of Alphafold 2 (https://github.com/YoshitakaMo/localcolabfold/releases/tag/v1.4.0), and used MMSeqs2 for the database search and clustering of protein sequences before performing the structural prediction (Mirdita et al. 2022). We used the whole ectodomains (see above) of the F-type and HSV/gB-type ENV proteins and predicted the trimer structures using the multimer mode of Alphafold 2 (AlphaFold2-multimer-v2) or Alphafold server (https://alphafoldserver.com/). We then visualised and characterised visible structures and disulfide bonds or cysteine bridges using Maestro (Schrödinger Release 2024-3) and grouped them by common substructures (Table S4). Structures of all predicted chromodomains in POL were confirmed by Alphafold. Ectodomains of viral class III fusogens of herpesvirus gB, rhabdovirus G, orthomyxovirus GP75/GP64-like proteins, baculovirus GP64, BEL/Pao-associated gB-like proteins were also included in this analysis when experimentally resolved structures were not available. Further, pairwise amino-acid identities were calculated within each viral family or errantivirus HSV/gB-type ENV group using curated ectodomain alignments. For each pairwise comparison, alignment columns containing a gap in either sequence were excluded, and identity was calculated as the number of identical residue–residue positions divided by the total number of compared residue–residue positions. These gap-excluded pairwise identities were used as descriptive measures of primary-sequence divergence and were not interpreted as evolutionary distances or molecular-clock estimates.

## Figure legends

**Supplementary Figure S1.**
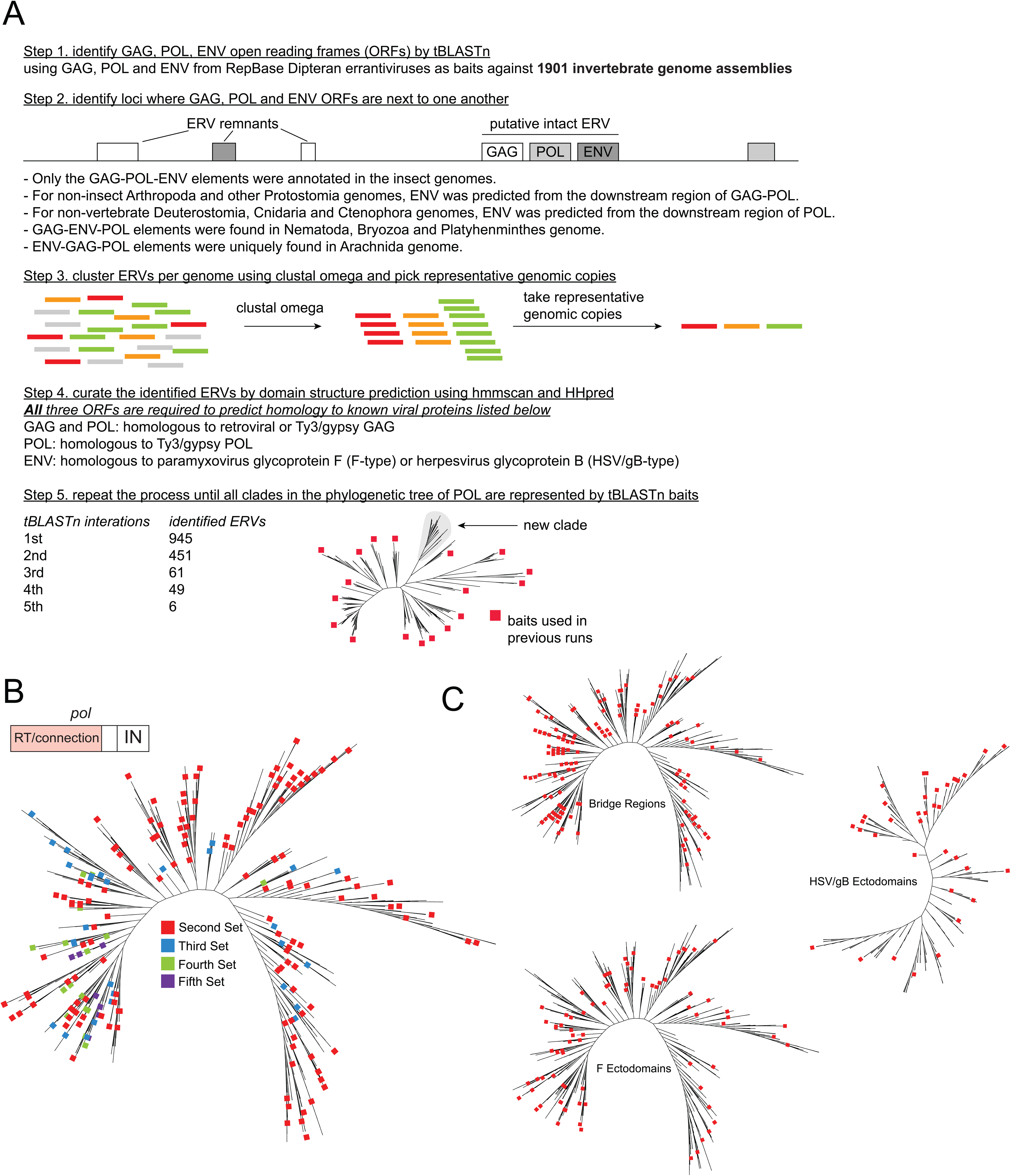
– Identification of errantiviruses in metazoan genomes. **(A)** Computational approach to identify and annotate *env*-containing retroelements in metazoan genomes. **(B)** A pruned maximum likelihood tree of POL extended RT/connection regions from errantiviruses with protein baits for consecutive tBLASTn searches annotated in colour, showing a comprehensive coverage of the baits in the tree. **(C)** POL extended RT/connection maximum-likelihood trees showing representative elements selected for AlphaFold2 structural analysis. Elements highlighted in red were used for structural prediction of the RNase H-containing bridge region, defined here as the canonical RNase H domain together with the C-terminal region between RNase H and Integrase. Representative elements used for AlphaFold2 analysis of F-type and HSV/gB-type ENV ectodomains are also indicated.

**Supplementary Figure S2.**
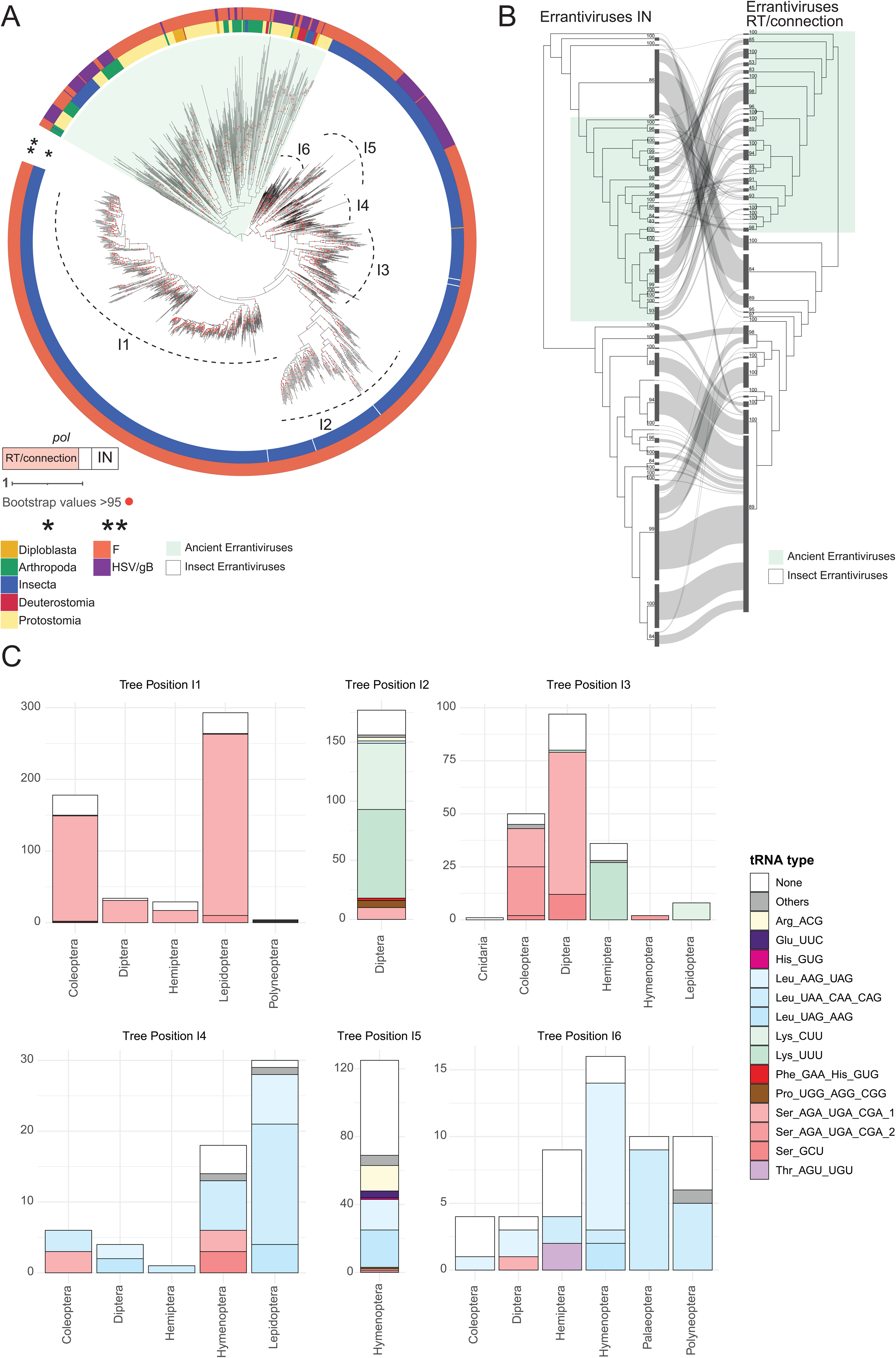
– Complete errantivirus tree and annotations of tRNA primer binding sites of insect errantiviruses. **(A)** Complete Maximum likelihood tree constructed on POL extended RT/connection regions of intact errantiviruses, with discrete phylogenetic groups within “insect errantiviruses” indicated (I1 – I6). “ancient errantivirus” group highlighted in green, “insect errantivirus group highlighted in white. Host phylogeny (*) annotated for insects (blue), non-insect arthropods (green), non-arthropod Protostomia (yellow), Deuterostomia (Red) and Diploblasta (Orange). ENV type (**) annotated for glycoprotein F (F) (orange), and glycoprotein B (HSV/gB) (purple). **(B)** A Sankey plot of POL extended RT/connection and IN maximum likelihood trees for errantiviruses, showing an overall phylogenetic congruence between them. **(C)** Shown are annotations of tRNA primer binding sites of “insect errantiviruses” within phylogenetic groups of POL extended RT/connection regions, demonstrating that the types of tRNAs used reflects the separation of POL extended RT/connection regions. Number of elements are indicated on Y axis. Sequence information of the tRNA PBS can be found in Table S1.

**Supplementary Figure S3.**
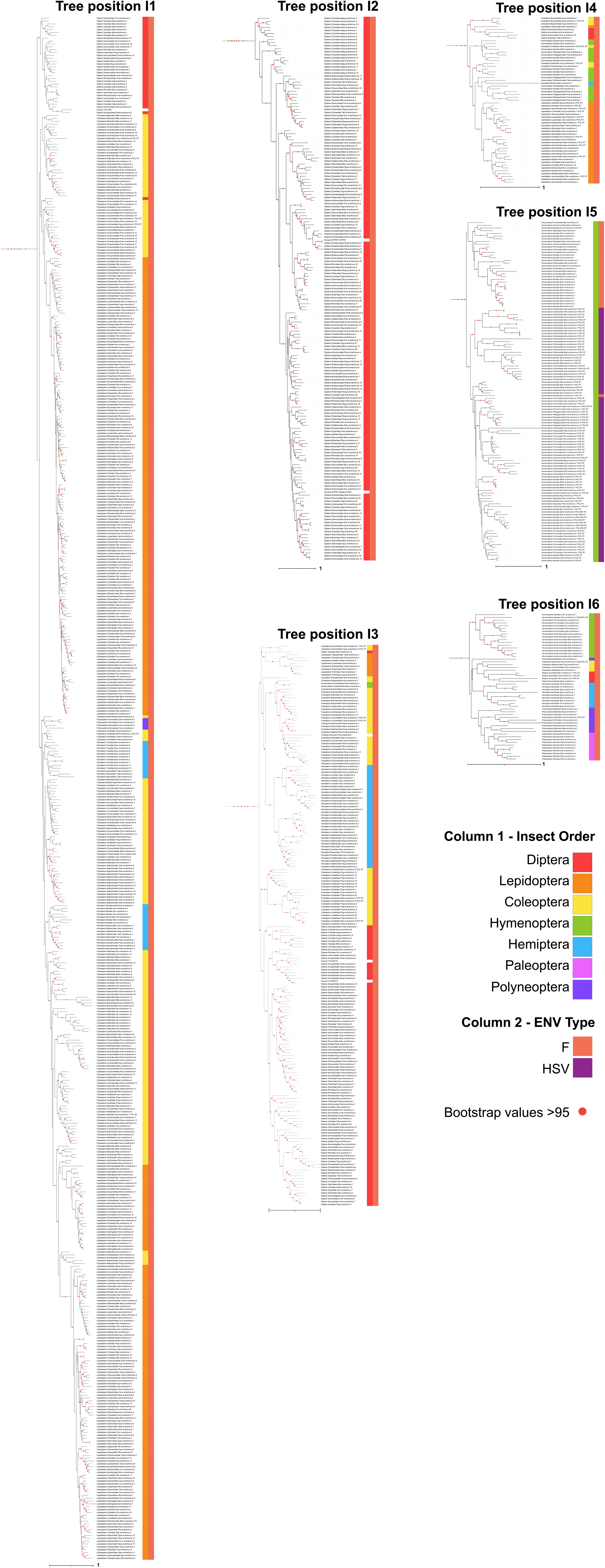
– Insect errantivirus clades. A maximum likelihood tree of POL extended RT/connection for the “insect errantivirus” group in isolation, with host phylogeny and *env* type, and discrete phylogenetic groups (I1-I6) indicated. Host orders (column 1) are annotated for Diptera (red), Lepidoptera (orange), Coleoptera (yellow), Hymenoptera (green), Hemiptera (cyan), Polyneoptera (pink) and Palaeoptera (purple). ENV types (column 2) are annotated for Glycoprotein F (F) (orange), and glycoprotein B (HSV) (purple). This figure provides the detailed tree underlying the insect portion of the representative POL phylogeny shown in the main figures and supports the conclusion that many insect errantivirus clades retain host-order or lower-level host-taxonomic structure.

**Supplementary Figure S4.**
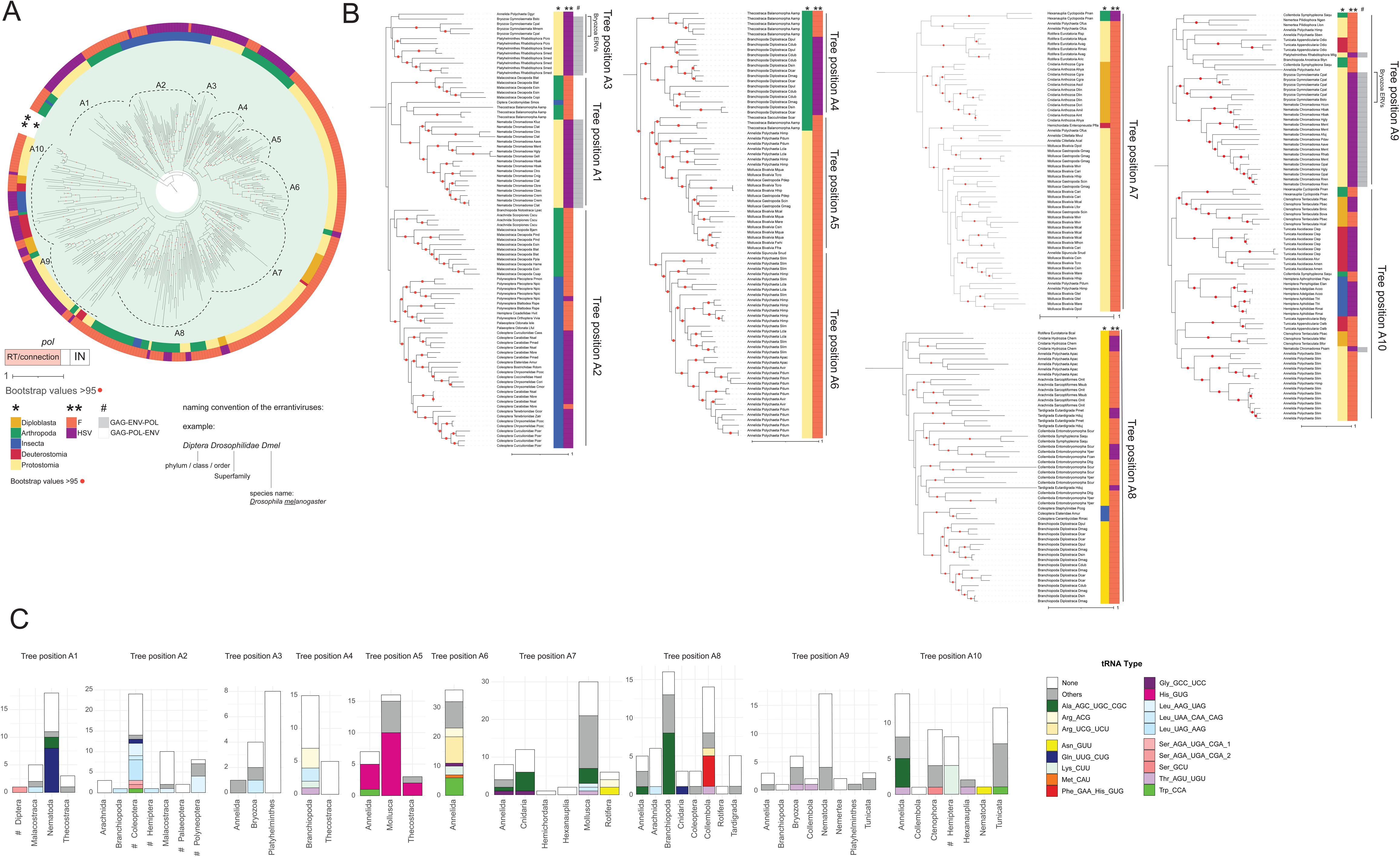
– Ancient errantivirus clades. **(A)** Maximum likelihood tree of POL extended RT/connection for the “ancient errantivirus” group in isolation, with host phylogeny and *env* type, and discrete phylogenetic groups (A1-A11) are indicated. Host phylogeny (*) is annotated for insects (blue), non-insect arthropods (green), non-arthropod Protostomia (yellow), Deuterostomia (Red) and Diploblasta (Orange). ENV type (**) annotated for glycoprotein F (F) (orange), and glycoprotein B (HSV) (purple). **(B)** Detailed view of phylogenetic groupings in “ancient errantiviruses” with host (*) and *env* (**) annotated. Elements with the non-canonical GAG-ENV-POL arrangement are indicated (#). Bryozoa elements with GAG-ENV-POL arrangement occurring in distinct positions of the tree are highlighted as an example of host-associated elements that do not form a single monophyletic group. Each element is named for the host species and simplified to Phylum/Class/Order, Superfamily, Genus and Species. For example, elements found in Diptera Drosophilidae *Drosophila melanogaster* are named as Diptera Drosophilidae Dmel. **(C)** tRNA annotations of “ancient errantiviruses” within phylogenetic groups of POL extended RT/connection. Insect orders in the groups A1, A2 and A7 are marked by hash. Number of elements indicated on Y axis. This figure provides the detailed phylogenetic context for the host-taxonomic clustering, ENV-type distribution and tRNA PBS patterns summarised in the main text.

**Supplementary Figure S5.**
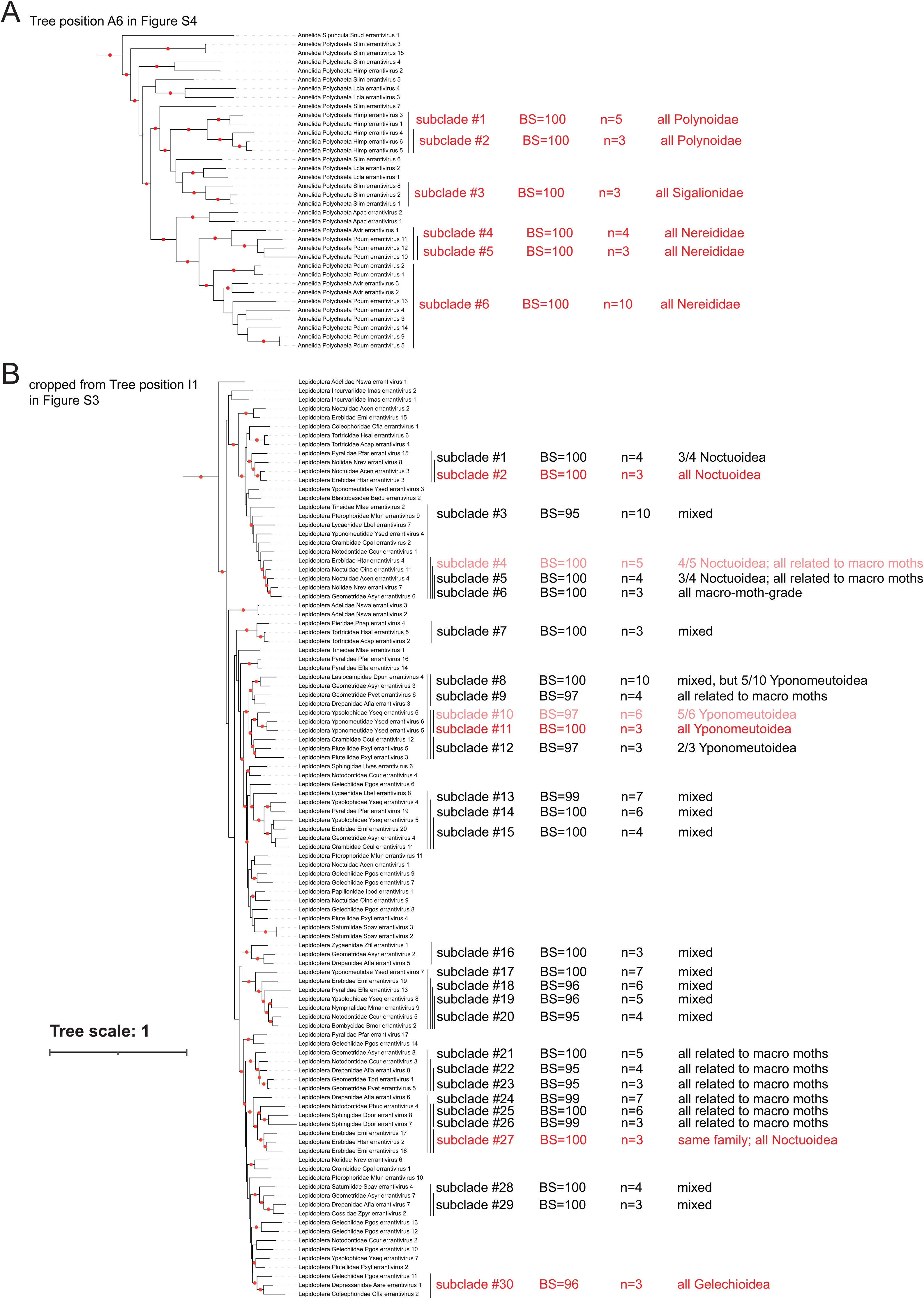
– Host-taxonomy concordance analyses of selected POL extended RT/connection subclades. Annelida-associated (A) and Lepidoptera-associated (B) errantivirus POL extended RT/connection subclades corresponding to tree position A6 (Fig. S4B) and I1(Fig. S3), respectively. Same tree scale for A and B. Highly supported internal subclades were identified using bootstrap support ≥95 and 3–10 descendant tips. Host-taxonomic concordance was assessed by comparing the observed distribution of host labels on the fixed POL topology with 10,000 random permutations preserving the observed label counts. In the Annelida clade, host-species labels were permuted and host-family labels were derived from the permuted species labels. Polynoidae, Sigalionidae and Nereididae are host families within Polychaeta. In the Lepidoptera clade, host-family labels were permuted and superfamily assignments were derived from the permuted family labels. Label “related to macro moths” refers to families represented in the tree from Drepanoidea, Geometroidea, Lasiocampoidea, Bombycoidea and Noctuoidea.Perfectly and mostly (≥80%) concordant subclades are highlighted in red and pink, respectively. These targeted analyses test whether host labels are non-randomly clustered on local POL trees, but do not estimate the number of horizontal transfer, duplication or loss events.

**Supplementary Figure S6.**
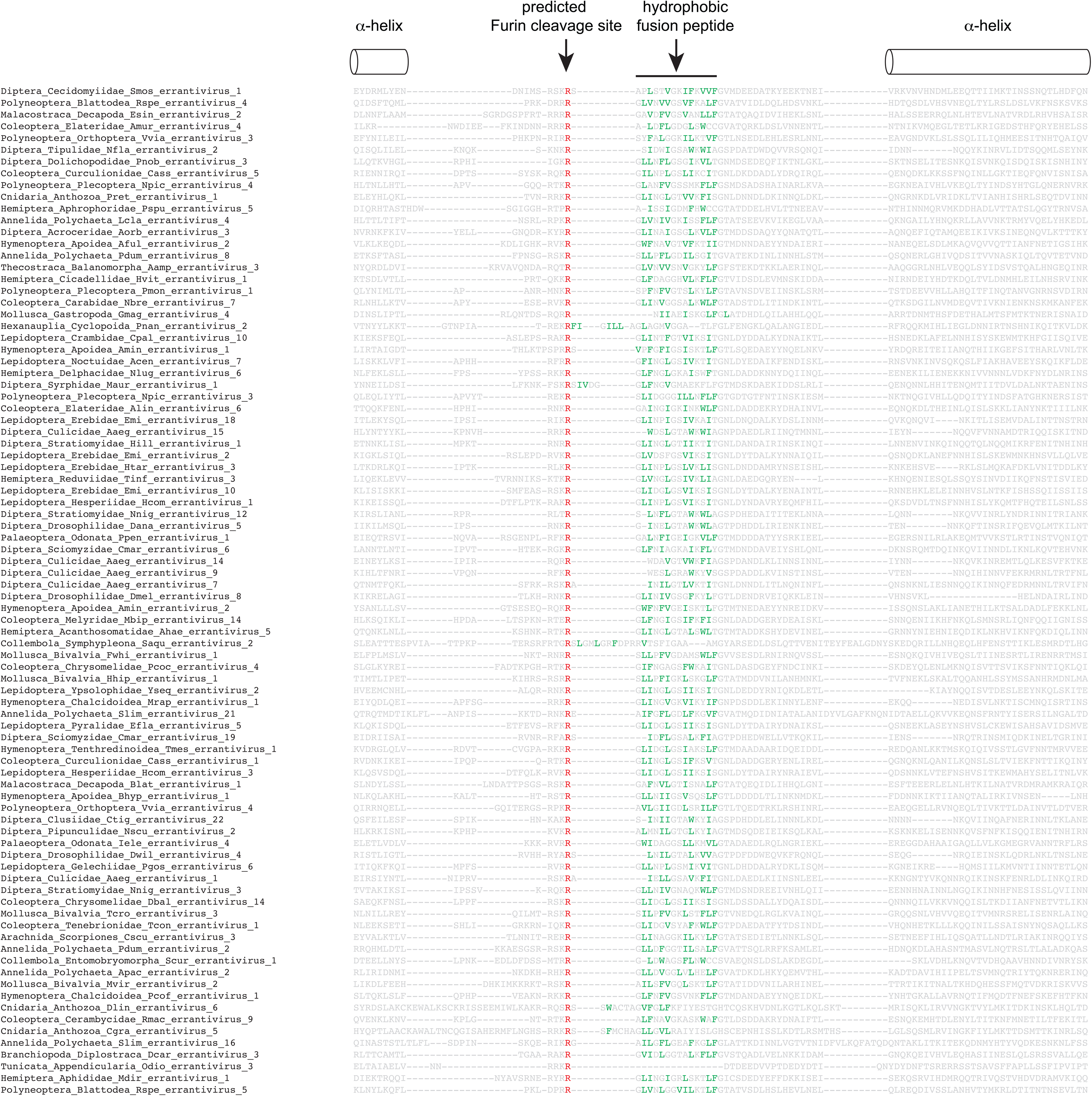
– Conserved predicted Furin cleavage sites and hydrophobic fusion peptides of glycoprotein F. Shown is a multiple sequence alignment of the middle part of the ectodomains of representative glycoprotein F. Arginine (R) residues in the putative Furin cleavage site and hydrophobic residues (Leu, Ile, Val, Phe, and Trp) in the fusion peptides are highlighted in red and green, respectively. Furin cleavage sites were predicted using ProP-1.0 (https://services.healthtech.dtu.dk/services/ProP-1.0/). Conserved α helix structures are shown upstream and downstream of the Furin cleavage site. This figure provides the alignment-level evidence supporting conservation of key F-type ENV motifs discussed in the main text.

**Supplementary Figure S7.**
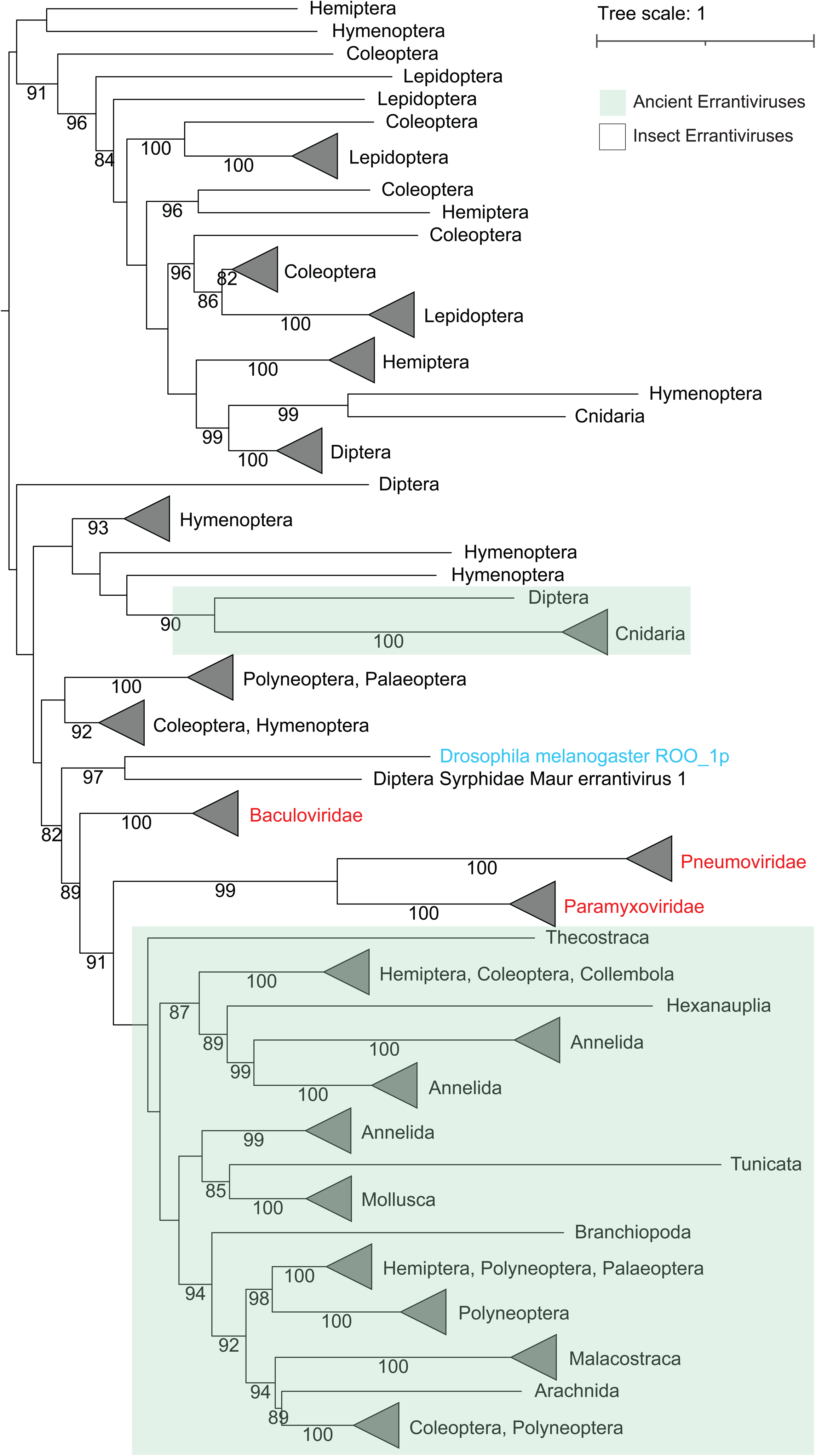
– Expanded phylogeny of F-type ENV ectodomains with viral and retroelement-associated homologues. Representative viral and non-*Ty3/gypsy* retroelement F-like proteins are included for comparison, including baculovirus F proteins, paramyxovirus and pneumovirus F proteins, and the *BEL/Pao*-associated Drosophila Roo F-like protein. The added viral sequences form family-level clades within the broader F-type ENV tree, while errantivirus F-type ENV proteins span a comparable scale of diversity.

**Supplementary Figure S8.**
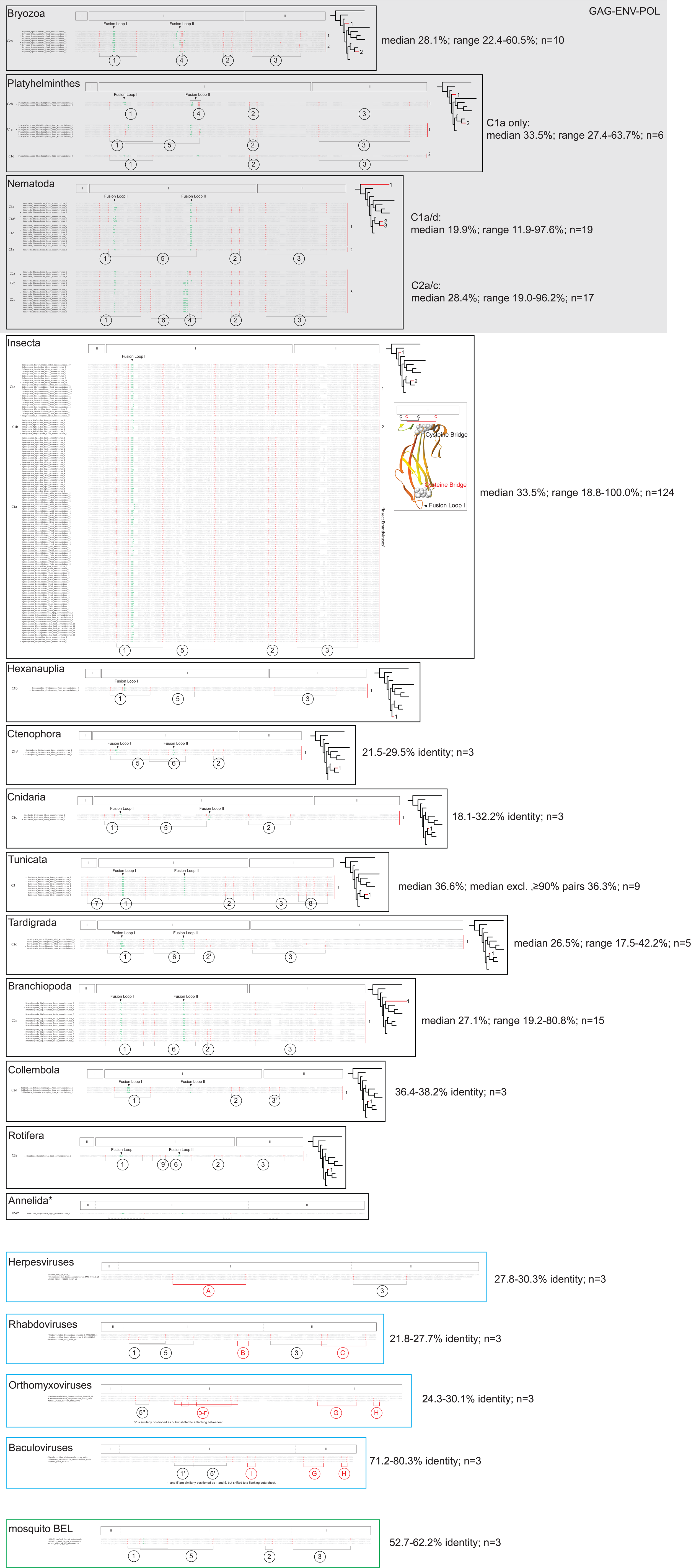
– Conservation of cysteine bridges in errantivirus HSV/gB ectodomains. Shown are alignments of domains I and II from errantivirus HSV/gB-type ENV ectodomains, together with representative viral and *BEL/Pao*-associated class III fusogens. Cysteine residues are highlighted in red and hydrophobic residues within the predicted fusion loop peptides are highlighted in green. Sequences from different metazoan orders, as well as representative viral or retroelement-associated groups, are aligned separately. Conserved cysteine bridges within and across the alignments are numbered, and ENV proteins with the GAG-ENV-POL arrangement are marked. Sequences used for structural prediction by Alphafold are indicated by asterisks, and those whose experimentally validated structures are available are marked by #. The structure of *Hymenoptera_Formicoidea_Tbic_errantivirus_1* is shown to highlight the position of the cysteine bridge in domain I of an insect errantivirus HSV/gB-type ENV ectodomain and the absence of fusion loop II. The single HSV/gB-type ENV identified in Annelida errantiviruses did not predict a trimeric structure and was therefore not included in the structural classification. For each viral family or errantivirus ENV group where there are at least three sequences, a summary of pairwise amino-acid identity is shown in the format “median x%; range y–z%; n = number of pairwise comparisons”. Raw data of pairwise amino-acid comparisons are also made available in Supplementary Table S2. This figure provides the alignment-level evidence that cysteine-bridge architectures are conserved within viral families and animal-associated errantivirus groups despite extensive primary-sequence divergence.

**Supplementary Figure S9.**
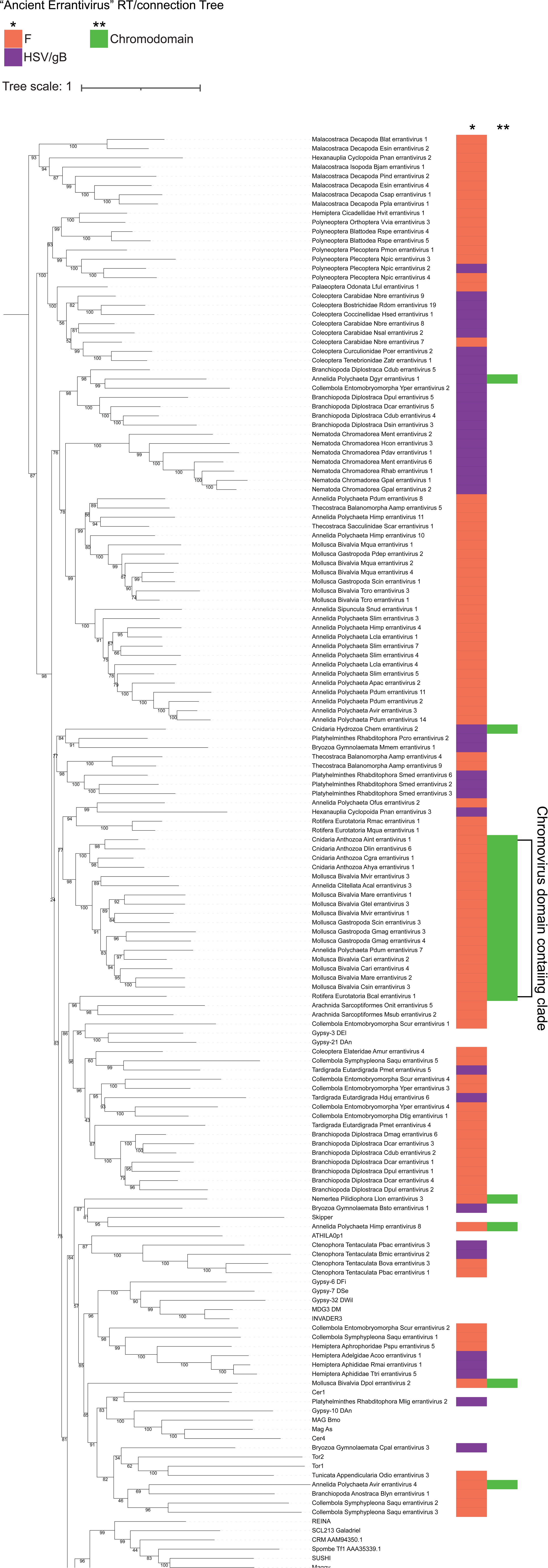
– Presence of chromodomain in “ancient” errantiviruses. – Maximum likelihood tree of POL extended RT/connection from representative errantiviruses along with known *Ty3/gypsy* elements. ENV type (*) annotated for Glycoprotein F (F) (orange), and glycoprotein B (HSV/gB) (purple). Chromodomain (**) annotated in purple.

**Supplementary Table S1. – Master errantivirus table.** Table of all 1512 newly identified *env-*containing errantiviruses identified in this study, with host phylogeny, genome assembly name and coordinates, contig size, type of ENV, copy number, tRNA PBS type, 3’ and 5’ LTR coordinates and PBS mismatches. The second tab summarises the tRNA PBS types identified in this study and their 12 nucleotides sequences.

**Supplementary Table S2. – Pairwise comparison of HSV/gB-type ENV sequences per subtypes.** Pairwise amino-acid identity comparisons among representative errantivirus HSV/gB-type ENV ectodomains and representative viral or retroelement-associated class III fusogens, including herpesvirus gB, rhabdovirus G, orthomyxovirus GP75/GP64-like proteins, baculovirus GP64 and BEL/Pao-associated gB-like proteins. Identities were calculated within each viral family or errantivirus ENV group over aligned residue–residue positions, excluding columns containing a gap in either sequence. The table includes the sequences analysed, all pairwise comparisons, and summary values used for figure labels, including sequence number, median identity, mean identity and identity range. C1, C2 and related labels refer to cysteine-bridge configurations defined in Figure 5 and Supplementary Figure S8. The values are used as descriptive measures of sequence divergence, not as molecular-clock estimates.

**Supplementary Table S3. – GAG-POL Ty3/gypsy elements identified in representative Porifera and Echinodermata genomes.** Table of full-length multicopy GAG-POL *Ty3/gypsy* elements identified in selected Porifera and Echinodermata genomes in which no intact GAG-POL-ENV errantiviruses were detected. The table includes host phylogeny, species name, genome assembly name and coordinates, contig size, copy number, predicted tRNA PBS type, 3’ and 5’ LTR coordinates, 12 nucleotide PBS coordinates, the corresponding tRNA/PBS sequences and PBS mismatches.

**Supplementary Table S4. – Structural Analysis of errantiviruses.** Lists of representative errantiviruses taken for structural analyses guided by Alphafold 2: RNase H domains found in the bridge region of *pol* (RNase H), F-type ENV ectodomain (F), HSV/gB-type ENV ectodomain, with cysteine bridge configuration indicated (HSV), and chromodomains found at the C-terminus of *pol* (chromodomain).

**Supplementary Table S5. – Node information for errantiviruses.** List of errantiviruses and phylogenetic node where they are found for errantivirus RT and IN (IN-RT Nodes Errantivirus Fig. S2) and RT and ENV from representative F-type errantiviruses (RT-ENV Nodes F-type Fig 4). Data was then used for congruence plots.

**Supplementary Table S6. – Genome assemblies analysed in this study.** List of genome assemblies with species name, size of assembly, class, family and number of tBLASTn hits for errantivirus ORFs. Genomes that we used to annotate errantiviruses are indicated. Tabs are individually detailed for Diptera, Lepidoptera, Coleoptera, Hymenoptera, Hemiptera, Polyneoptera, Palaeoptera, non-insect Arthropoda, non-arthropod Protostomia, Cnidaria, Ctenophora, Porifera, and non-vertebrate Deuterostomia.

## Acknowledgements

We acknowledge Julius Brennecke, Kirsten Senti and Alex Greenwood for valuable comments in preparation of the manuscript. This work was supported by the Australian Research Council (DP210102385).

## Author contributions

S.C. and R.H. analysed and interpreted the data and wrote the manuscript. The idea of this work was conceived by R.H., and S.C. and R.H. together made key findings.

## Data Availability

The following data are available from the git repository (https://github.com/RippeiHayashi/errantiviruses_2025/tree/main): the code used to annotate and analyse genomic copies of errantiviruses, genomic sequences and open reading frames of the newly identified errantiviruses, pdb files for the predicted structures of ENV proteins, the bridge region and the C-terminal half of POL, sequences of GAG, POL and ENV open reading frames that are used for tBLASTn search, multiple sequence alignments and the tree files that are generated in this study for the structural and phylogenetic analyses.

